# Anaerobic isoprene reduction by *Pelotomaculum* sp. From *Eucalyptus*-leaf sediments and its impacts on methanogenesis

**DOI:** 10.64898/2026.02.19.706888

**Authors:** Samikshya Giri, Zoe I. Kelley, Anna E. Rockwood, Sabrina Beckmann

**Affiliations:** Department of Microbiology, Oklahoma State University, OK 74078, USA

**Author notes:** Author Contributions Samikshya Giri and Dr. Sabrina Beckmann contributed to the design and implementation of the research, analysis of the results, and manuscript writing. Dr. Sabrina Beckmann conceived the original idea and supervised the project. Anna E. Rockwood and Zoey Ilene Kelley assisted with experimental work and data collection. Conflicts of Interest The authors declare no conflicts of interest.

**Keywords:** Isoprene reduction, *Pelotomaculum* sp, methyl-butenes, methanogenesis

## Abstract

Isoprene is a ubiquitous biogenic volatile organic compound (VOC) with atmospheric concentrations comparable to those of methane. Its high reactivity in the atmosphere significantly influences methane concentrations, contributing to adverse effects on the climate. Understanding the role of isoprene and its link to methane metabolism is crucial to addressing climate change. The fate of isoprene and its potential microbial degraders in soil, particularly in anaerobic environments, remains poorly investigated, underscoring the need for comprehensive studies.

Our study provides physiological evidence for anaerobic microbial isoprene reduction and its influence on methanogenesis in *Eucalyptus*-leaf sediment. In our anaerobic culture-based studies, we have demonstrated that isoprene is reduced to three products, 3-methyl-1-butene, 2-methyl-1-butene, and 2-methyl-2-butene. Our findings suggest that *Pelotomaculum* sp. is capable of anaerobic isoprene reduction, as indicated by 16S rRNA analysis and dilution-to-extinction studies. Anaerobic microbial isoprene reduction simultaneously inhibited methane formation. Methanogenesis was completely inhibited in microcosm cultures amended solely with isoprene. In cultures supplemented with isoprene and carbon sources, only hydrogenotrophic methanogenesis was inhibited, indicating competition for hydrogen between the methanogens and isoprene degraders. Using a culture-based approach and molecular analysis, this study provides novel insights into the microbial dynamics of anaerobic isoprene reduction and the interplay between isoprene reducers and methanogens.

**Graphical Abstract:** 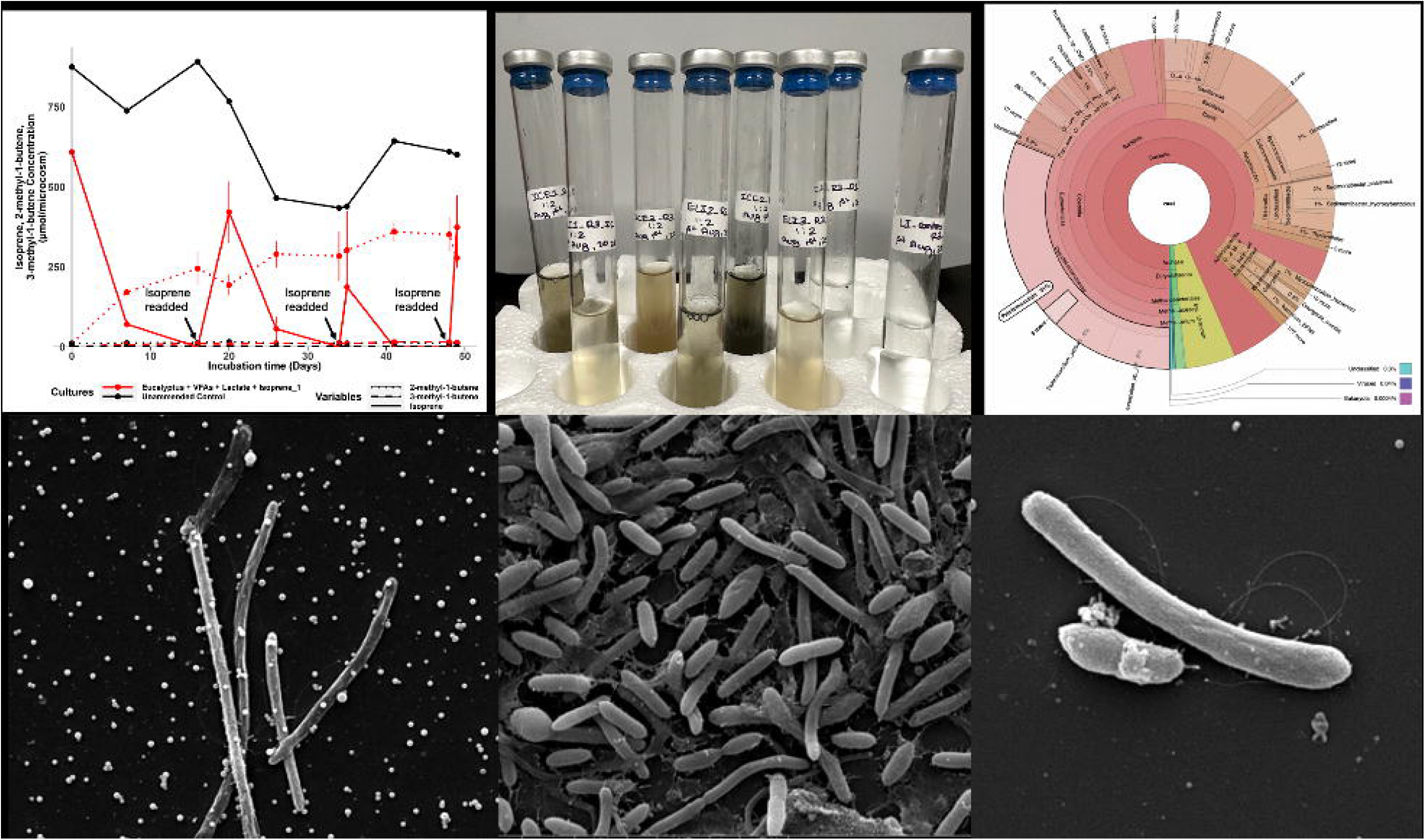

## Introduction

Isoprene (2-methyl-1,3-butadiene) is a fundamental precursor to various biogenic and anthropogenic compounds and is synthesized ubiquitously by diverse organisms, including plants (Sharkey *et al*., 2007), humans (Gelmont *et al*., 1981), and select bacteria (Fall & Copley, 2000). It is among the most volatile organic compounds, with a boiling point of 34°C, and is highly reactive due to its carbon-carbon double bonds (Carrión *et al*., 2020). In plants and bacteria, isoprene is thought to function as a signaling molecule that aids in stress protection, as evidenced by its increased production under stress conditions, such as exposure to hydrogen peroxide, high temperature, and high salinity (McGenity *et al*., 2018; Xue & Ahring, 2011). Approximately 90% of the total isoprene emitted into the atmosphere originates from plants as a byproduct of photosynthesis (Sharkey *et al*., 2007; Pacifico *et al*., 2009; Lahr *et al*., 2015). Isoprene was typically the dominant volatile organic compound emitted by Eucalyptus species, accounting for 64–100% of total emissions (He *et al*., 2000). Previous studies have shown that high levels of NOx and VOCs emitted from the canopies of these plants have detrimental effects on air quality and human health (Ashworth *et al*., 2013).

The atmospheric concentration of isoprene is comparable to that of methane, with global emissions reaching up to 500 Tg per year (Carrión *et al*., 2020). Studies indicate that isoprene prolongs methane’s atmospheric residence time by consuming radicals such as hydroxyl (OH^-^), nitrate (NO ^-^), and ozone (O_3_) that would otherwise react with methane (Charlson *et al*., 1992; Pacifico *et al*., 2009; Murrell *et al*., 2020). At high concentrations, isoprene reacts with nitric oxide (NO) to form nitrogen oxides, which are harmful to human health and air quality (Pacifico *et al*., 2009; Ervens *et al*., 2011; Wyche *et al*., 2014). Secondary aerosols formed by the photooxidation of isoprene are also thought to contribute to cooling the Earth’s atmosphere by absorbing and scattering solar radiation (McGenity *et al*., 2018). The net effect of isoprene on global warming remains unclear; however, unraveling the sources and potential sinks of isoprene in the environment, such as soil, is critical for understanding the biogeochemical cycle of isoprene and its role in climate change.

Studies of aerobic isoprene-degrading microorganisms have established soil as a critical sink for isoprene (Cleveland & Yavitt, 1997; Gray *et al*., 2015; McGenity *et al*., 2018). Aerobic catabolism of isoprene has been detected in several *Rhodococcus* species and in novel *Nocardioides, Ramlibacter,* and *Variovorax* strains that use isoprene as a carbon and energy source (Carrión *et al*., 2020). Several studies on isoprene-degrading soil microorganisms have established *Rhodococcus* sp. AD45 as a model organism for aerobic isoprene metabolism (Crombie *et al*., 2015; Carrión *et al*., 2020; Sims *et al*., 2022). Sims *et al*. (2022) purified and characterized isoprene monooxygenase (*IsoMO*), a four-component multi-enzyme complex that catalyzes the first step of isoprene degradation by converting isoprene to 3,4-epoxy-3-methyl-1-butene. Genomic analyses of Rhodococcus sp. AD45 and other isoprene-degrading microorganisms have led to the discovery of the isoA gene, which encodes the large alpha subunit of *IsoMO* and serves as a biological marker for isoprene metabolism in aerobic microbes (Khawand *et al*., 2016).

Considerable progress has been made in our understanding of the aerobic microbial metabolism of isoprene (Dawson *et al*., 2023), but research on the anaerobic metabolism of isoprene is limited to *Acetobacterium wieringae*. Studies by Kronen *et al*. (2019, 2023) and Jin *et al*. (2022) demonstrated that isoprene serves as an electron acceptor in *Acetobacterium wieringae*, being converted to methyl butenes with no further conversion observed. To investigate the distribution of anaerobic isoprene reducers within *Acetobacterium* species, Kronen *et al*. (2019) tested pure isolates of *A. woodii* (DSM 1030), *A. malicum* (DSM 4132), and *A. wieringae* (DSM 1911) and found isoprene reduction activity limited to *A. wieringae* strain ISORED-2. Similarly, Jin *et al*. (2022) detected this activity in *A. wieringae* strain Y. Both studies collected samples from wastewater treatment plants.

Our study aims to expand understanding of isoprene-reduction activity among anaerobic microorganisms, thereby increasing the diversity of known isoprene-reducing anaerobes. Previous studies on isoprene-reducing anoxic cultures have reported that isoprene inhibits methanogenesis, but this effect has not been thoroughly investigated (Kronen *et al*., 2019; Jin *et al*., 2022). Our research addresses these two critical knowledge gaps. First, we provide robust physiological and molecular evidence for anaerobic microbial isoprene reduction through culture-based experiments, 16S rRNA analyses, and metagenomic analyses. We identify *Pelotomaculum* sp. as a putative anaerobe that reduces isoprene to methyl-butenes. Second, we establish a relationship between methanogenesis and anaerobic isoprene reduction by leveraging GC and 16S rRNA analyses. By identifying *Pelotomaculum* sp. as a putative anaerobic isoprene reducer and elucidating the relationship between isoprene reducers and methanogens, our study provides valuable insights into the nuanced anaerobic microbial interactions that shape atmospheric chemistry and climate dynamics.

Our findings underscore the complex role of anaerobic microbial communities in modulating isoprene and methane dynamics, highlighting the need for continued exploration of their effects on atmospheric composition and climate change.

## Materials and Methods

### Enrichment cultures

The Isoprene 99% stock solution (Sigma Aldrich), 3-methyl-1-butene ≥ 95.0% (TCI), 2-methyl-2-butene ≥ 99.0% (Thermo Scientific), and 2-methyl-1-butene ≥ 98% (Thermo Scientific) were obtained and stored at-4°C. Helium (>99.9999% purity), nitrogen gas (>99.99% purity), N2/CO2 (25% CO2, 75% N2), and air (zero-grade purity) were obtained from Airgas Gas Radnor, PA, USA. Standard solutions were prepared in 120 ml flasks using 70 ml of anaerobic minimal medium. Isoprene was added to the stock solution using a gas-tight glass syringe to prepare the standards. 3-methyl-1-butene, 2-methyl-2-butene, and 2-methyl-1-butene were added to their respective stock solutions using a gas-tight glass syringe in separate flasks as standards.

Anaerobic minimal medium for sample inoculum was prepared using MgCl2 x 6H2O (0.4 g l-1), KBr (0.09 g l-1), KCl (0.66 g l-1), and CaCl2 x 2H2O (0.1 g l-1). The medium was dispensed into 120 ml culture flasks, crimp-sealed with Teflon-faced rubber septa, autoclaved, and degassed with N2. The headspace of the culture bottles was filled with N2/CO2 (80:20, v/v). NaHCO3 (1M, 30 ml l-1), NH4Cl (5 g l-1) / KH2PO4 (4 g l-1), 30 ml l-1 mixture; trace element mixture (1 ml, Nitriloacetic acid, 1.5 g l-1; MgSO4 x 7H2O, 3 g l-1; NaCl, 1 g l-1; CaCl2 x 2H2O, 0.1 g l-1; KAI(SO4)2 x 12 H2O, 0.02 g l-1; Na2SeO3 x 5H2O, 0.3 mg l-1; Na2WO4 x 2H2O, 0.4 mg l-1; FeSO4 x 7H2O, 0.1 g l-1; MnSO4 x H2O, 0.5 g l-1; CoSO4 x 6H2O, 0.18 g l-1; CuSO4 x 5H2O, 0.01 g l-1; NiCl2 x 6H2O, 0.03 g l-1; Na2MoO4, 0.01 g l-1; ZnSO4 x 7H2O, 0.18 g l-1; H3BO3, g l-1; distilled water, 1 l); 1 ml vitamin solution (Biotin, 2 mg l-1; thiamine hydrochloride, 5 mg l-1; pyridoxine hydrochloride, 10 mg l-1; folic acid, 2 mg l-1; riboflavin, 5 mg l-1; lipoic acid, 5 mg l-1; D-Ca-pantothenate, 5 mg l-1; Vitamin B12, 0.1 mg l-1; 4-aminobenzoic acid, 5 mg l-1; nicotinic acid, 5 mg l-1; distilled water, 1 l). The vitamins and trace element solutions were filter-sterilized, and all other solutions were autoclaved.

Soil detritus was collected from beneath Eucalyptus trees at a forested site in California, USA, from the surface soil (5 cm depth) containing decomposed leaf litter. Soil sediment was collected aseptically with sterile spatulas and transferred into sterile containers. Samples were transported to the laboratory on ice and kept at 4°C until further processing. This material served as the inoculum for enrichment cultures and subsequent DNA extraction. Dry soil sediment (1.3 g) was inoculated into culture bottles containing 70 ml of sulfate-free anaerobic minimal media (Widdel & Bak, 1992). A mixture of volatile fatty acids (VFAs), including acetate, butyrate, propionate, succinate, and formate at 10 mM, along with L/D lactate (10 mM), was added to the media. Isoprene was added as an electron acceptor from a 99.9% stock solution using a 100 μl gas-tight glass syringe, yielding a final concentration of (100-300) μmol in the liquid media. Dilution-to-extinction series were conducted under the same culture conditions using 25 ml Belco tubes with 5 ml of anaerobic minimal media and 5 ml of enriched culture. Controls included a dead control and an unamended control. The dead control was prepared by supplementing the culture with 0.5 g l-1 sodium azide. All incubations were performed in triplicate. Culture samples for molecular analyses were promptly frozen at-80°C. Methane, hydrogen, and acetate concentrations were monitored over 12 months using regular GC analyses before replenishing the media, substrates, and isoprene.

### Isoprene, methylbutenes, and methane analysis

Isoprene, methylbutenes, and methane were analyzed every 2-3 weeks by directly collecting 100 μl of gas from the culture headspace. Separation of isoprene and methylbutenes was achieved on a GasPro PLOT column (60 m × 0.32 mm, Agilent Technologies) with nitrogen as the carrier gas at 5 mL/min. Detection was performed with a flame ionization detector (FID). The GC oven was programmed with an initial hold at 50 °C for 30 s, followed by a ramp of 20 °C/min to a final temperature of 250 °C. Manual injection of the collected headspace samples ensured consistent analysis of substrate and product concentrations. Gas samples (100 μl) were extracted from the flask using a pressure-lockable gas-tight syringe (Fisher Scientific, UK) and manually injected into the GC.

### DNA extraction and Illumina sequencing

DNA was extracted from 1 ml culture using a FastDNA^TM^ Spin Kit for Soil (MP, USA) according to the manufacturer’s instructions. The extracted DNA was stored at-80°C until further use. Amplification of the V4 region of the 16S rRNA gene was amplified by PCR with the modified universal primers 515F and 806R, which target bacteria and archaea (Caporaso *et al*., 2011). The 515F primer was modified by introducing a barcode (Parada *et al*., 2016). PCR products were purified with AMPure XP (Beckman Coulter) following the Illumina MiSeq protocol. Samples were then sequenced on an Illumina MiSeq Sequencer (Illumina, USA) using paired-end 16S community sequencing at the Microbial Analysis, Resources, and Services Center for Open Research Resources and Equipment at the University of Connecticut. The resulting 16S rRNA gene amplicon sequences were analyzed in Mothur (Schloss PD *et al*., 2009) at the Pete Supercomputer Facility, Oklahoma State University. Stability files derived from the raw data were used to generate contigs. Anomalies and duplicate sequences were excluded. Forward and reverse sequences that passed default quality control were merged, while non-overlapping sequences were discarded. The resulting output file was aligned with the SILVA nr_v138_1. Align sequence files (SILVA 2023). Poorly aligned sequences were discarded, and overhangs at both ends of the aligned sequences were removed. Sequences differing by only 2 bp were merged into a single sequence to reduce the impact of sequencing errors. A thorough chimera search was conducted to detect and exclude chimeric sequences from further analysis. Lineage classification was then applied, focusing on bacterial and archaeal lineages classified to the genus level, whereas eukaryotic and unidentified lineages were excluded. Finally, taxa with abundances below 1% were excluded to refine the analysis and enhance clarity. 16S rRNA amplicon sequencing data used for community analysis were deposited in the NCBI Sequence Read Archive (SRA) under BioProject PRJNA1153434. Sequence data for the samples are available under BioSample SAMN43393446.

### Shotgun metagenomic sequencing and bioinformatic analysis

Purified DNA samples were sent to Novogene Co., Ltd. (Beijing, China) for shotgun metagenomic sequencing. Sequencing libraries were prepared by randomly fragmenting genomic DNA, followed by end-repair, A-tailing, adapter ligation, size selection (∼350 bp inserts), and PCR amplification. Libraries were quantified with Qubit and qPCR, assessed for size distribution on a fragment analyzer, and sequenced on the Illumina NovaSeq X Plus platform to generate 150 bp paired-end reads. Raw reads were processed with fastp v0.23.1 to remove adapter contamination and reads with >10% ambiguous bases or >50% low-quality bases. Host DNA sequences were filtered with Bowtie2 v2.2.4. High-quality reads were assembled de novo with MEGAHIT v1.2.9 (meta-large preset), and contigs <500 bp were discarded. Open reading frames (ORFs) were predicted on assembled scaffolds with MetaGeneMark v2.1; ORFs <100 aa were excluded. Predicted ORFs were clustered with CD-HIT v4.5.8 at 95% identity and 90% coverage to construct a non-redundant gene catalog (unigenes). Gene abundance was estimated by mapping clean reads back to the catalog with Bowtie2. Taxonomic annotation was performed by aligning unigenes against the MicroNR database with DIAMOND v2.1.6 (blastp, e-value ≤ 1e−5). Functional annotation was performed using DIAMOND against the KEGG, eggNOG, CAZy, VFDB, and PHI databases. Antibiotic resistance genes were identified by comparison with the CARD database, and mobile genetic elements were annotated using ISfinder, Integrall, and Plasmid databases. Downstream statistical analyses, including α-diversity indices, PCA, PCoA, NMDS, clustering, and correlation analyses, were conducted in R (v2.15.3–3.0.3) using packages such as corrplot, ggplot2, vegan, and VennDiagram. Taxonomic profiles were visualized using Krona.

### Microscopic analyses

Scanning Electron Microscopy (SEM) and Fluorescence Microscopy were used to evaluate cell quantity and morphology. For SEM, samples were fixed with 2% glutaraldehyde and stored at 4°C. Samples were centrifuged at 14,000 × g for 15 min, and the supernatant was discarded. A small amount of the supernatant was retained in the tube to prevent the cell pellet from drying completely. To coat the sample onto glass slides, coverslips were rinsed with methanol, then with deionized (DI) H_2_O. The coverslip was evenly coated with a 1 µg ml-1 poly-lysine solution. After drying, the coverslips were rinsed with deionized (DI) H_2_O. 50 µl of the fixed cell suspension was applied to a poly-lysine-coated coverslip. After drying, the coverslip with the sample was incubated in a standard aldehyde fixative for 15 min to cross-link poly-lysine with the sample. The coverslip was then thoroughly rinsed with DI H2O to remove excess fixative. The samples were dehydrated through a series of ethanol concentrations (50%, 70%, 90%, 95%, 100%) for 15 min each. The samples were then washed twice for 5 min each with HMDS and coated with Au-Pd. The coated samples were scanned and imaged using an FEI Quanta 600 FEG Environmental Scanning Electron Microscope (Oklahoma State University). Using fluorescence microscopy, cells were stained with SybrGreen I (Invitrogen, USA), and images were captured with a BX51 epifluorescence microscope (Olympus, USA) as described by Lunau *et al*. (2005).

## Results

### Anaerobic reduction of isoprene in cultures with volatile fatty acids and lactate as carbon sources

Microcosms enriched with *Eucalyptus*-leaf detritus and supplemented with volatile fatty acids (VFAs) and lactate as carbon and energy sources, and isoprene as an electron acceptor, were incubated for 1 year, then replenished with isoprene, carbon sources, and media (50%). After replenishment, isoprene, its reduced products (2-methyl-1-butene, 3-methyl-1-butene, 2-methyl-2-butene), and methane were quantified by gas chromatography (GC, Fig. 1, FigS.1).

**Fig. 1.**
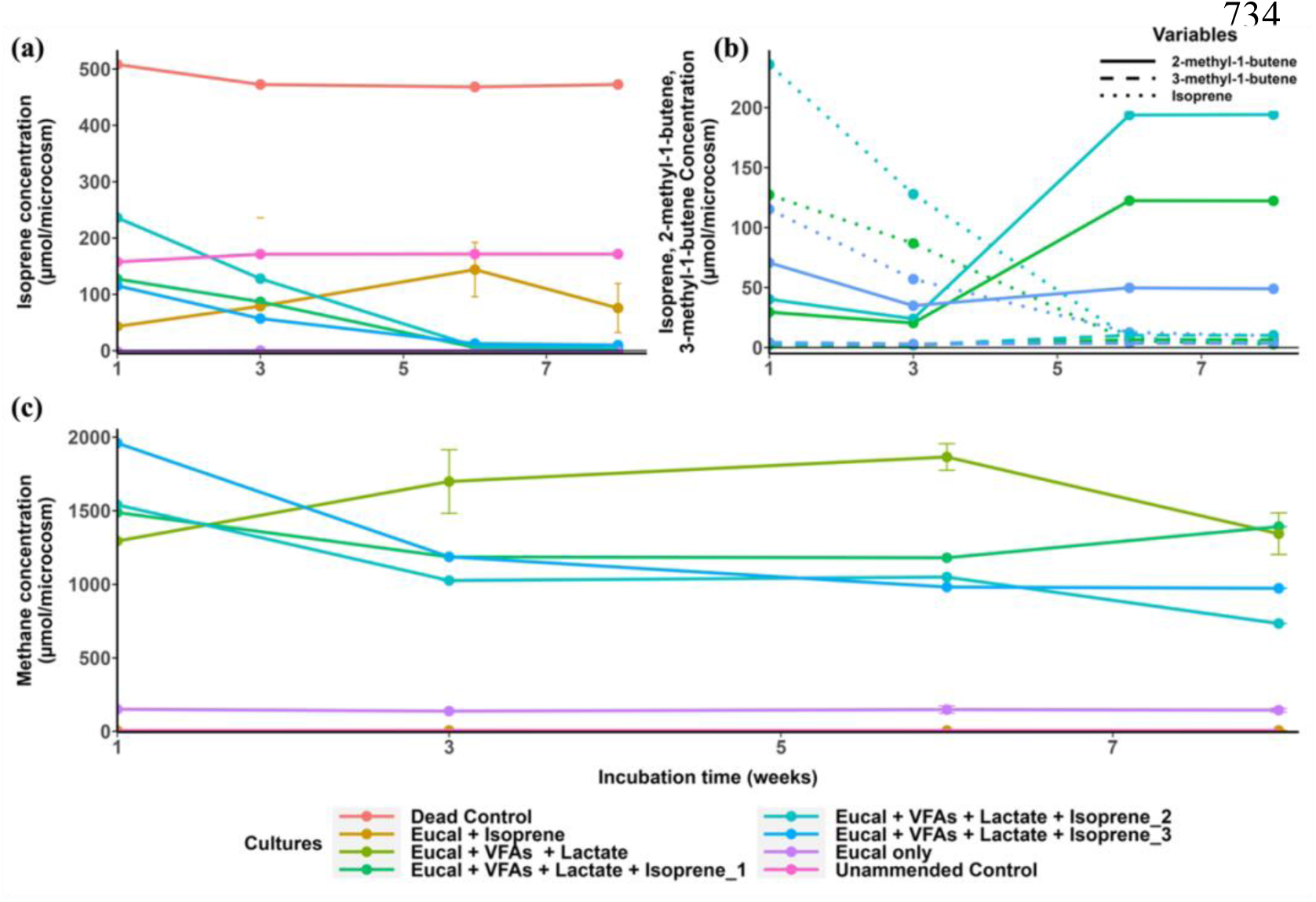
Concentrations of isoprene, methylbutenes, and methane in anaerobic *Eucalyptus*-leaf cultures after one year of incubation and replenishment of substrates, media, and isoprene. GC analyses began 1 week after replenishment and incubation. (**a**) Depletion of isoprene. **(b**) Depletion of isoprene and production of methylbutenes (2-methyl-1-butene and 3-methyl-1-butene) were observed in microcosms containing isoprene and carbon sources (VFAs and lactate). **(c)** Methane production occurred in all microcosms. Error bars indicate one standard deviation (n = 3).

In the first round of enrichment cultures, 159.49 ± 66.49 µmol of isoprene per microcosm was fully depleted within 6 weeks (Fig. 1 a). Concurrently, 121.80 ± 72.64 µmol/microcosm of 2-methyl-1-butene (∼97%) and 6.77 ± 3.3 µmol/microcosm of 3-methyl-1-butene (∼3%) were produced (Fig. 2 b). Trace amounts of 2-methyl-2-butene appeared inconsistently in select cultures. These occurrences remained near the detection limit and were therefore considered negligible for the purposes of quantitative comparison. Isoprene was partially reduced and methyl butenes were generated in all replicates with *Eucalyptus-*leaf detritus inoculum and isoprene but no carbon sources (Eucal + Isoprene). By Week 8, isoprene levels in the *Eucalyptus*-leaf cultures amended with isoprene averaged 75.74 ± 43.74 µmol/microcosm with concomitant reduction into the major product 2-methyl-1-butene (60.74 ± 35.07 µmol/microcosm), the minor product 3-methyl-1-butene (4.12 ± 2.38 µmol/microcosm; FigS2a). 2-methyl-2-butene was not detected in any replicates of the *Eucalyptus*-leaf cultures amended with isoprene. No isoprene reduction products were observed in the dead control or unamended control, although small isoprene depletion (6.99%) was noted in the dead control after 60 days. Self-degradation of isoprene in anaerobic cultures after 45 days has been previously reported (Kronen *et al*., 2019). In the control group with only the *Eucalyptus*-leaf inoculum (Eucal only), neither isoprene nor its reduced products were observed.

**Fig. 2.**
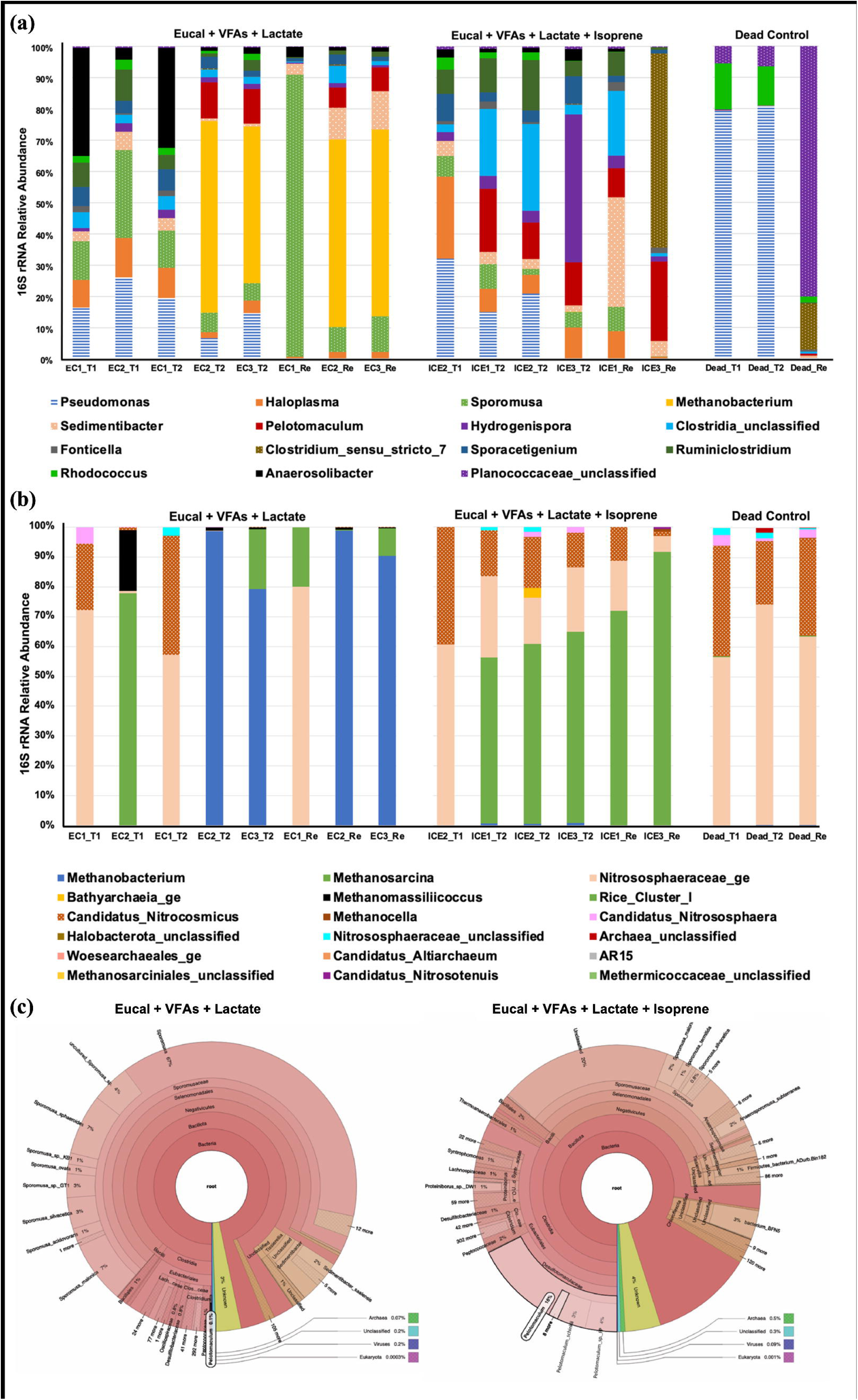
Composition of the bacterial and archaeal populations, classified at the genus level, in *Eucalyptus*-leaf detritus enrichments amended with VFAs and lactate as carbon sources and isoprene as an electron acceptor, at three timepoints: T1 (2 months), T2 (12 months), and ReT1 (1 week after readdition of substrates, media, and isoprene). Microcosms were inoculated with *Eucalyptus*-leaf sediment and incubated for a year before replenishing the culture. (a) Bacterial community composition. (b) Archaeal community composition. (c) Krona representation of microbial abundance in microcosms without (i) and with added isoprene (ii) as an electron acceptor.

Isoprene reduction persisted through three cycles of medium replenishment (50%), isoprene addition, and carbon-source supplementation. Reduced isoprene products were consistently observed in dilution-to-extinction cultures, indicating that isoprene reduction was not transient. Methylbutenes were not further reduced, consistent with previous findings (Kronen *et al*., 2019; Jin *et al.,* 2022). Interestingly, methanogenic activity differed across all microcosms.

### Methane formation in microcosms is dependent on isoprene reduction

Methane formation (methanogenesis) was evident in all *Eucalyptus*-leaf microcosms except those without a carbon source amendment (Eucal + Isoprene; Fig. 1c). The highest methanogenic activity was observed in *Eucalyptus*-leaf cultures amended with VFAs and lactate (n=2), at 1343.62 ± 199.4 µmol/microcosm, followed by cultures amended with VFAs, lactate, and isoprene (n=3), at 1032.78 ± 332.32 µmol/microcosm, indicating partial inhibition of methanogenesis by isoprene. Residual methanogenesis, 143.82 ± 13.13 µmol/microcosm, was observed in *Eucalyptu*s-leaf cultures without carbon supplementation (FigS2b). No methanogenic activity was detected in any of the 3 replicates of *Eucalyptus*-leaf enrichments amended with only isoprene, the uninoculated control, or the dead control. These results suggest that isoprene inhibits methanogenesis, likely by competing with methanogens for residual organic substrates, such as formate or hydrogen (H₂), which isoprene reducers could use as electron donors.

### Bacterial and archaeal community of VFA and lactate-driven anaerobic isoprene reduction

To explore bacterial and archaeal community succession during enrichment, 16S rRNA gene amplicon community analysis was performed on all *Eucalyptus*-leaf isoprene enrichment cultures before and during isoprene reduction (Fig. 2, FigS3a). Enrichment cultures revealed significant shifts in microbial dynamics, with *Pseudomonas* dominating at 15.09% during the first 12 months of incubation, followed by *Haloplasma* at 6.01% across all replicates (Fig. 2a). Notably, a rapid transition in the microbial community occurred within a week of replenishing the medium (50%), isoprene, and carbon sources (VFAs and lactate), displacing *Pseudomonas* with other genera. In cultures amended with isoprene and carbon sources, hydrogen-producing anaerobes of the genera *Clostridium* (*Ruminoclostridium*, *Clostridium sensu stricto_7*, *Clostridia unclassified*) and *Hydrogeniospora* were predominant (Fig. 2a). Although their precise role in isoprene transformation remains unclear, they likely supply hydrogen to anaerobes engaged in isoprene reduction. Beyond *Clostridium*, *Pelotomaculum* emerged as a key microorganism in isoprene-amended cultures. A comparative analysis of Krona representations shows that in cultures without isoprene, *Pelotomaculum* accounted for only 0.1% of the overall microbial community (Fig. 2a). In contrast, in isoprene-supplemented cultures, the relative abundance of *Pelotomaculum* increased to 18%.

*Methanobacterium* was identified as the fourth most abundant microbial genus, with an overall relative abundance of 4.12% across all culture replicates (Fig. 2b). Methane production was detected in all cultures supplemented with carbon sources, regardless of the presence of isoprene (Fig. 1c). However, *Methanobacterium* was the only methanogen among the top 15 most abundant taxa and the most dominant microbe in cultures without isoprene (Fig. 2a), except in 1 of 3 replicates of the *Eucalyptus*-leaf cultures amended with VFAs and lactate (EC1; Fig. 2a), which did not produce any methane. 16S rRNA analysis revealed the absence of methanogens throughout the enrichment period. The 16S rRNA profile of EC1 was similar to that of the microbial community of the *Eucalyptus*-leaf cultures amended with only isoprene (FigS2) and the dead controls (Fig. 2a). EC1 was dominated by *Sporomusa*, a homoacetogen with an exceptionally low hydrogen (H₂) threshold (Laura & Jo, 2023). The predominance of *Sporomusa* in EC1, coupled with the absence of methane production, suggests that EC1 had an insufficient hydrogen pool to support the growth of the hydrogenotrophic methanogen *Methanobacterium*. 16S rRNA analysis of the archaeal community indicated the presence of methanogens in both cultures with and without isoprene (Fig. 2b). However, distinct methanogenic populations dominated under these conditions: *Methanosarcina* was the predominant methanogen in cultures with isoprene, whereas *Methanobacterium* dominated in cultures without isoprene. This suggests a shift in methanogenic community composition driven by isoprene reduction, with *Methanosarcina* potentially outcompeting *Methanobacterium* in isoprene-enriched environments.

### Different archaeal community composition in the *Eucalyptus*-leaf microcosms *vs Eucalyptus*-leaf microcosms amended with isoprene

Bacterial 16S rRNA analysis revealed no significant differences in the top 20 most abundant microbial genera between *Eucalyptus*-leaf isoprene-amended cultures and *Eucalyptus*-leaf-only enrichments (FigS3b). However, archaeal community composition differed markedly between the two enrichment groups (Fig. 3). Methanogenic species were absent in all three replicates of the *Eucalyptus*-leaf isoprene-amended cultures, consistent with GC analysis (Fig. 1c), which showed no methane production in those cultures. These cultures were dominated by *Nitrososphaeraceae_ge* and *Candidatus Nitrocosmicus.* In contrast, *Eucalyptus*-leaf cultures lacking isoprene harbored methanogens spanning three metabolic types: methylotrophic (*Bathyarchaeia_ge, Methanomassiliicoccus*), hydrogenotrophic (*Rice Cluster I, Methanocella*), and versatile methanogens such as *Methanosarcina*, capable of hydrogenotrophic, methylotrophic, and acetoclastic methanogenesis. These shifts suggest that competition between methanogens and isoprene-reducing microorganisms, driven by limited hydrogen and carbon availability, constrains methanogenesis in isoprene-amended cultures.

**Fig. 3.**
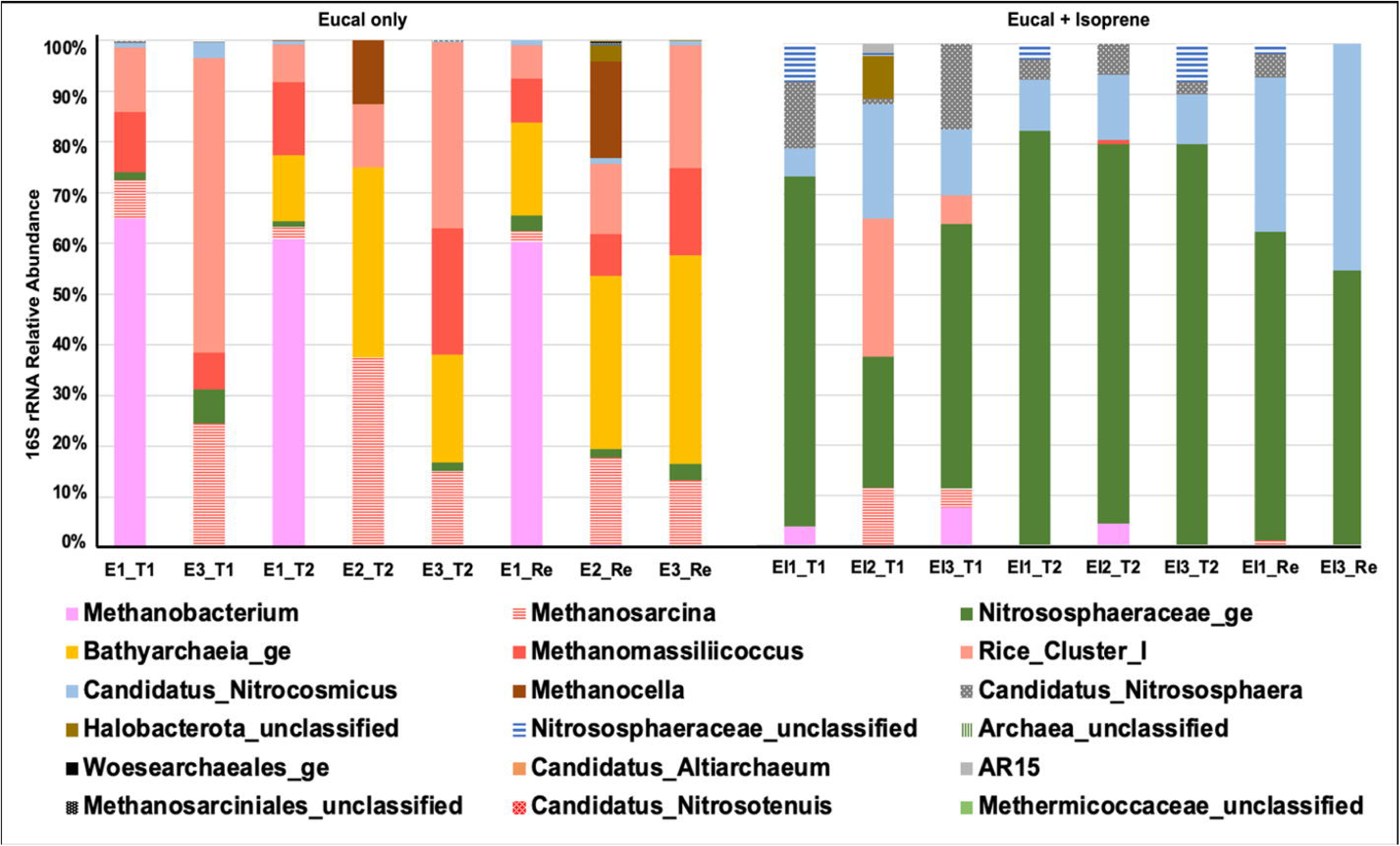
Archaeal community composition at the genus level in Eucalyptus-leaf sediment microcosms with and without isoprene at three timepoints: T1 (2 months), T2 (12 months), and ReT1 (1 week after readdition of substrates, media, and isoprene).

### Dilution-to-extinction isolation of isoprene-reducing microorganism(s)

A dilution-to-extinction isolation study is underway to isolate isoprene-reducing microorganisms from isoprene-amended and carbon-source-amended cultures (VFAs and lactate; Fig. 4). Fluorescence and scanning electron microscopy images of the original enrichments (Fig. 4. a-g) and the first round of dilution cultures (Fig. 4b-c) provided insights into the succession of the microbial community structure. The original enrichment was obscured by soil debris, limiting visualization of cell morphology. The dilution enriched two distinct microbial cell types: rod-shaped and coccoid microorganisms (Fig. 4. d-g).

**Fig. 4.**
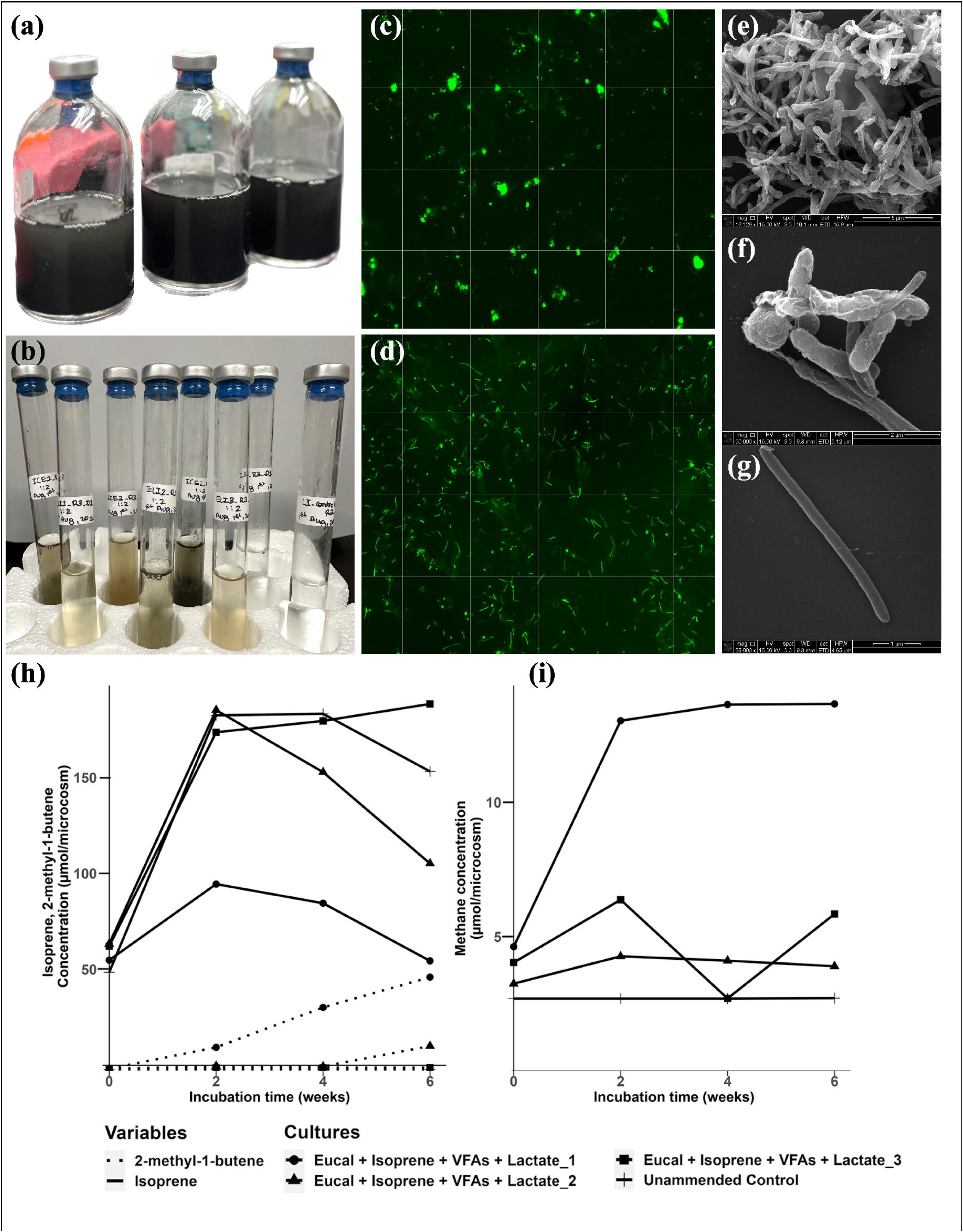
Dilution-to-extinction cultures of Eucalyptus-leaf sediment amended with VFAs and lactate as carbon and energy sources, and isoprene as an electron acceptor. (a) Original Eucalyptus-leaf microcosms after 3 rounds of media and substrate supplementation (Eucalyptus+VFAs+Lactate) and isoprene addition (Eucal+VFAs+Lactate_1, Eucal+VFAs+Lactate_2). (b) First dilution culture, Eucal+VFAs+Lactate_1. (c-d) Fluorescence microscopy images of the 1^st^ dilution culture, Eucal+VFAs+Lactate_1, after 6 weeks of incubation. (e-g) Scanning Electron Microscopy (SEM) images of the 1^st^ dilution culture, Eucal+VFAs+Lactate_1. (h) Shows the generation of 2-methyl-1-butene and the depletion of isoprene in the 1^st^ dilution-to-extinction culture. (i) Depicts methane formation in the 1^st^ dilution-to-extinction culture.

One replicate (n=3) of the *Eucalyptus*-leaf culture dilution series amended with VFAs and isoprene produced 45.96 µmol/microcosm of 2-methyl-1-butene, a key product of isoprene reduction, within the first 6 weeks (Fig. 4h). This culture also exhibited the highest methanogenic activity, yielding 13.67 µmol of methane (Fig. 4i). Eucalyptus-leaf cultures amended with isoprene, VFAs, and lactate overcame the initial lag phase after 6 weeks, reduced isoprene to 10.02 µmol/microcosm of 2-methyl-1-butene, and produced 3.90 µmol of methane. Neither methane nor 2-methyl-1-butene was detected in any of the unamended controls.

In the second round of dilution-to-extinction of *Eucalyptus*-leaf cultures amended with isoprene and carbon sources (VFAs and lactate; n=3), one replicate (Fig. 4 h-i) reduced isoprene from 115.46 µmol/microcosm to 101.33 µmol of 2-methyl-1-butene/microcosm, but no methane production was observed. The second replicate reduced isoprene from 128.98 µmol to 116.93 µmol of 2-methyl-1-butene/microcosm, and no methane formation was observed either. *Eucalyptus*-leaf cultures enriched with VFAs and lactate as carbon sources showed consistent isoprene reduction throughout the dilution cultures, whereas those supplemented with lactate only showed no isoprene reduction activity in the dilution-to-extinction cultures. These findings underscore the impact of dilution on the microbial community (Fig.S4), particularly the replacement of methanogens by isoprene reducers, highlighting the potential competitive disadvantage methanogens face relative to isoprene-degrading microorganisms, even in the presence of preferred carbon sources like VFAs and lactate.

### Enrichment of *Pelotomaculum* in the dilution-to-extinction isolation cultures

Microbial relative-abundance analysis revealed substantial enrichment of *Pelotomaculum*, increasing from 18% in the original culture to 30% and 31% in the two dilution-to-extinction cultures (Fig. 5). Sequences similar to *Pelotomaculum schinkii* and *Pelotomaculum sp.* accounted for 6% of the total microbial community in both enrichment replicates, while archaeal species comprised only 1% of the overall microbial community. *Methanosarcinales* comprised 6% of the total archaeal abundance and 0.08% of total microbial abundance. These findings suggest that isoprene potential confers a competitive advantage to *Pelotomaculum species*, supporting their role in energy conservation through isoprene reduction.

**Fig. 5.**
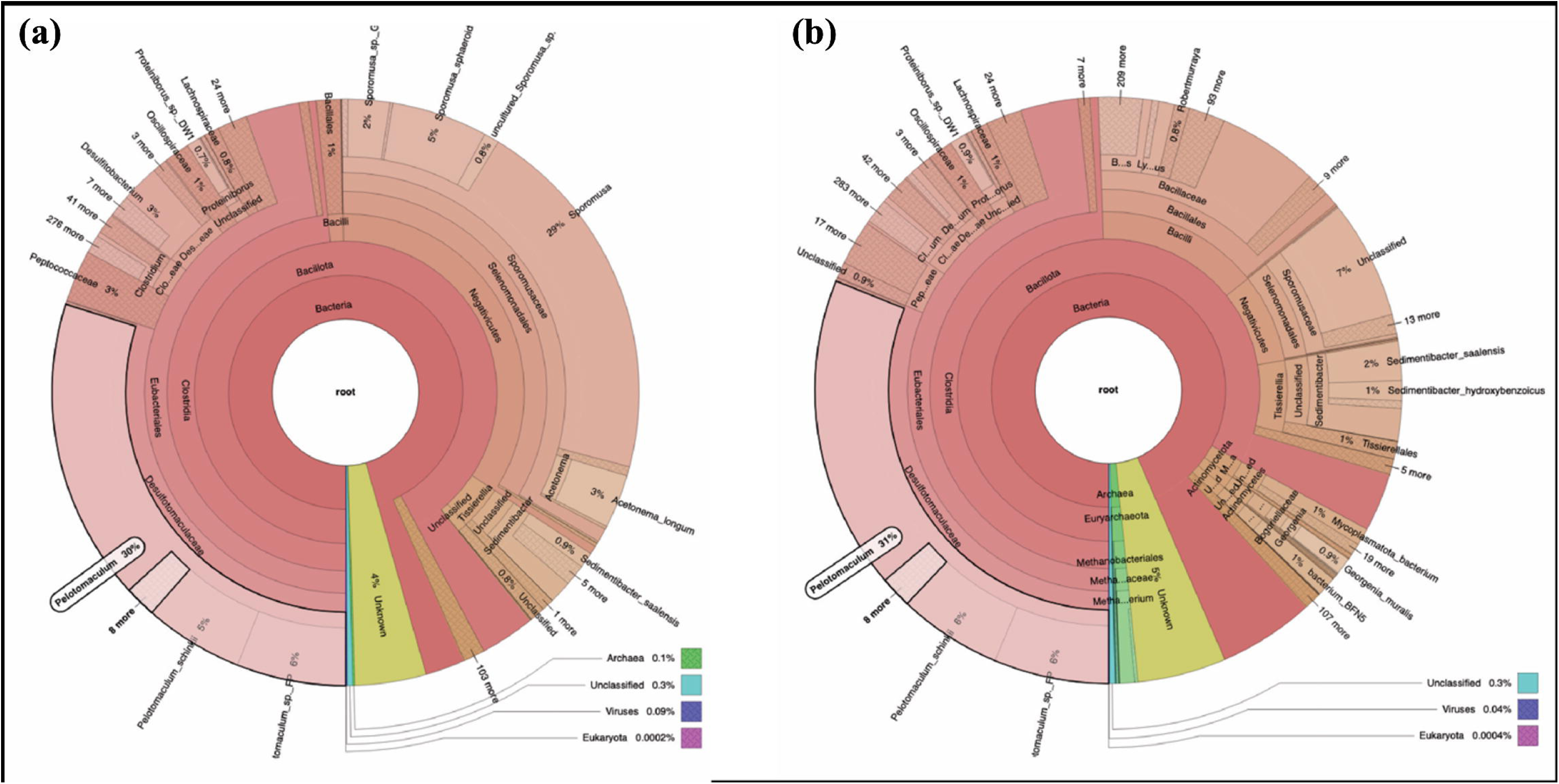
Krona representation of *Pelotomaculum* relative abundance in the 3^rd^ dilution-to-extinction cultures. (a) Eucal + Isoprene + VFAs + Lactate_1 and (b) Eucal + Isoprene + VFAs + Lactate_2.

### Long-term enrichment cultures amended with isoprene, volatile fatty acids (VFAs), and lactate

To assess whether isoprene-reduction activity could be sustained over multiple replenishments of isoprene after depletion (Fig.S6), *Eucalyptus*-leaf enrichments amended with isoprene, VFAs, and lactate from the 7th round of dilution cultures (ICE1_D7 and ICE2_D7) were monitored for 45–50 days, with periodic re-amendments of isoprene (Fig. 6). In both cultures, rapid and repeated depletion of isoprene followed each addition, accompanied by the formation of the reduced products 2-methyl-1-butene and 3-methyl-1-butene. In enrichment ICE1_D7, isoprene concentrations declined from 636.8 µmol/microcosm at day 0 to 308.7 µmol/microcosm by day 7 and further to 46.0 µmol/microcosm by day 34 (Fig. 6a). Correspondingly, 2-methyl-1-butene and 3-methyl-1-butene increased from negligible levels at time 0 to 180.6 µmol/microcosm and 7.8 µmol/microcosm, respectively, by day 34 (Fig. 6a). After each re-addition of isoprene, similar depletion trends were observed, with isoprene decreasing from ∼346.6 µmol/microcosm to <50 µmol/microcosm within one week, indicating sustained metabolic activity. In ICE2_D7, isoprene decreased sharply from 1374.5 µmol/microcosm at day 0 to 210.3 µmol/microcosm by day 6, representing an 85% reduction, while 2-methyl-1-butene and 3-methyl-1-butene accumulated to 195.3 µmol/microcosm and 8.9 µmol/microcosm, respectively (Fig. 6b). After three consecutive re-additions of isoprene (day 20, 28, and 35), isoprene was again rapidly consumed, reaching < 3 µmol/microcosm by day 30 and complete depletion by day 35, with sustained accumulation of reduced products up to 465 µmol/microcosm (2-methyl-1-butene) and 26.7 µmol/microcosm (3-methyl-1-butene; Fig. 6b).

**Fig. 6.**
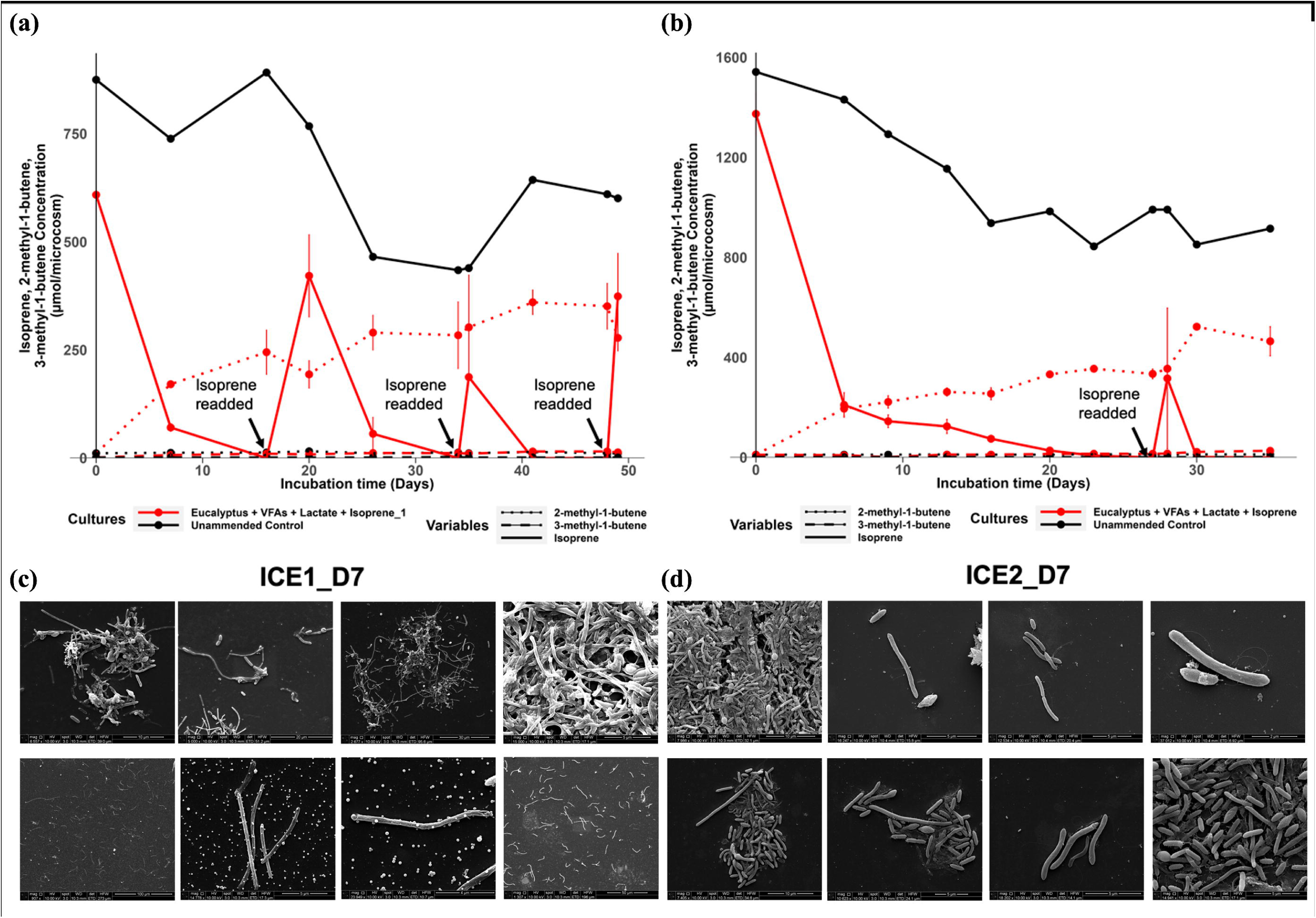
Time-course profiles of isoprene and its reduced products in long-term enrichment cultures of (a) ICE1_D7 and (b) ICE2_D7 in the 7^th^ round of dilution, and (c) Scanning Electron Micrographs of enrichment cultures ICE2_D7 replicates after 22 days of incubation, and (d) ICE2_D7 at 41 days of incubation. Replicates of both enrichments, ICE1_D7 and ICE2_D7, were supplemented with *Eucalyptus*-leaf sediment as inoculum, volatile fatty acids (VFAs), lactate, and isoprene. The unamended control was amended with the same carbon sources and isoprene as the amended control but lacked the inoculum. Isoprene was periodically reamended (dark arrows), and the concentrations of isoprene (solid lines), 2-methyl-1-butene (dotted lines), and 3-methyl-1-butene (dashed lines) were monitored. Error bars represent standard deviation (n = 4).

Both reduced products of isoprene were below detection in the unamended controls, even though isoprene concentration declined slightly throughout the incubation, confirming that the reduction of isoprene to 2-methyl-1-butene and 3-methyl-1-butene was biologically mediated. Across both enrichments, isoprene consumption rates remained constant after each supplementation, indicating that the microbial community retained its reductive capacity over extended incubation. The reproducible product profiles and stable conversion efficiency highlight a metabolically active, sustained isoprene-reducing consortium under VFA- and lactate-supplemented anaerobic conditions.

### Scanning electron micrographs of the long-term enrichment cultures at different growth stages

Scanning electron microscopy (SEM) of *Eucalyptus*-leaf cultures amended with isoprene, VFAs, and lactate (ICE2_D7 and ICE1_D7 enrichments) revealed a morphologically diverse microbial community after 22 and 41 days of incubation, respectively (Fig. 6c,d).

After 21 days of incubation, the micrographs showed numerous elongated rod-shaped and filamentous cells, typically 1–5 µm in length, arranged in dense clusters and surface-associated aggregates (Fig. 6c). Individual rods displayed smooth surfaces and occasional curvature, with some cells appearing in pairs or short chains, suggestive of active division. The abundance of filamentous forms and compact cell groupings indicates the potential establishment of a syntrophic network characteristic of cooperative anaerobic metabolism. These structural arrangements likely facilitate close physical contact between metabolic partners. The presence of a continuous extracellular matrix and well-organized aggregates suggests a metabolically active consortium in log phase, coinciding with the period of maximal isoprene depletion observed in the headspace measurements, as indicated by GC analyses (Fig. 6a).

After 41 days of incubation, Scanning Electron Microscopy (SEM) imaging of the Eucalyptus-leaf cultures amended with isoprene, VFAs, and lactate (ICE1_D7; Fig. 6a) revealed distinct morphological and structural differences compared with the earlier 22-day culture (Fig. 6a). The microbial community appeared less densely packed, with more dispersed, thinner filamentous cells distributed across the field of view. Long, slender filaments measuring 5–10 µm and individual rods were interspersed among small aggregates, suggesting partial disintegration of previously compact cell clusters (Fig. 6d). At higher magnification, cells exhibited smoother surfaces and less extracellular matrix material than in earlier stages, consistent with a decline in active biofilm formation and a transition to a more quiescent physiological state. The presence of residual filamentous networks indicates persistence of syntrophic structures, though at lower density, reflecting a shift from peak metabolic activity associated with log phase toward a maintenance phase following substrate depletion. Overall, these morphological changes suggest that the consortium remained viable but metabolically less active, indicating resource depletion during the later stages of incubation, i.e., the stationary phase.

### Anaerobic microbial transformation of isoprene with only lactate as a carbon source

*Eucalyptus*-leaf cultures amended with isoprene and lactate converted isoprene to methyl butenes within 6 weeks across all replicates (n=3). The primary products of isoprene reduction were 335.93 ± 38.32 µmol of 2-methyl-1-butene (∼97%) and 16.39 ± 3.73 µmol of 3-methyl-1-butene (∼3%). No isoprene depletion or reduced products of isoprene were observed in the dead controls, whereas minimal isoprene depletion (12.8%) occurred in the unamended controls. Reduced products of isoprene were not detected in either the dead controls or the unamended controls, confirming that abiotic processes did not contribute to isoprene reduction. Methanogenesis was observed in all culture replicates except the controls. After 8 weeks, the *Eucalyptus*-leaf cultures amended with isoprene and lactate produced 320.68 ± 114.53 µmol of methane. In contrast, *Eucalyptus*-leaf cultures amended with lactate but not isoprene produced 473.8 ± 88.03 µmol, suggesting that isoprene reduction partially inhibits methanogenesis. In the dilution-to-extinction cultures, solely lactate-amended microcosms exhibited no detectable isoprene reduction or methanogenic activity, unlike those amended with VFAs and lactate.

## Discussion

### Isoprene reduction driven by *Pelotomaculum* species in the microbial community

The genus *Pelotomaculum* comprises strictly anaerobic, Gram-positive, rod-shaped, spore-forming syntrophic bacteria (Morris *et al*., 2013). They are uniquely adapted to environments lacking traditional electron acceptors, using low-energy substrates such as propionate, butyrate, lactate, and ethanol, often in partnership with methanogens (De Bok *et al*., 2005; Westerholm *et al*., 2021). Because of their reliance on low-energy substrates, *Pelotomaculum* species are slow-growing and considered part of the ‘rare biosphere’, contributing only a small fraction to the overall microbial community (Tveit *et al.,* 2015). Among these substrates, propionate oxidation is the least energetically favorable, with a standard Gibbs free energy change (ΔG°) of +73.8 kJ/mol (Thauer *et al*., 1977; Hanselmann, 1991; Amend & Shock, 2001; Hidalgo-Ahumada *et al*., 2018), as shown in Equation (1). Despite being thermodynamically unfavorable, *Pelotomaculum schinkii* can persist under these energy-limited conditions by forming a syntrophic relationship with methanogens. The overall free energy change when coupled with methanogenesis is approximately −15 kJ/mol (Hidalgo-Ahumada *et al*., 2018), making the process energetically feasible, as shown in Equation (3). This syntrophic interaction is essential for *P*. schinkii’s survival in anaerobic environments with minimal energy availability.

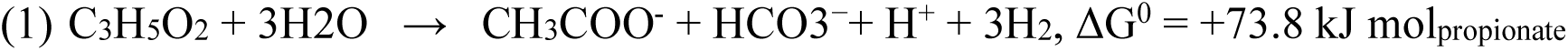

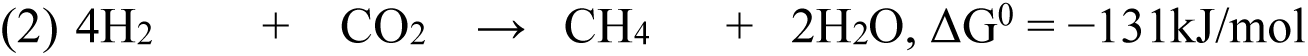

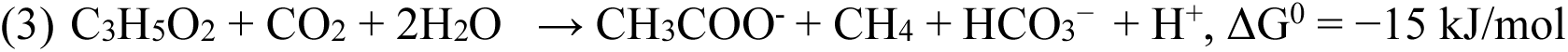

By using alternative electron acceptors such as isoprene, *Pelotomaculum* can overcome the thermodynamic constraints of propionate oxidation without a syntrophic methanogenic partner. Theoretically, *P. schinkii* can couple propionate oxidation with isoprene reduction, yielding a standard Gibbs free energy of −316 kJ/mol (Kronen *et al*., 2023), as shown in Equation (4). This suggests that using isoprene as an electron acceptor could be a more favorable metabolic strategy for *P. schinkii* than its traditional syntrophic partnership with methanogens.

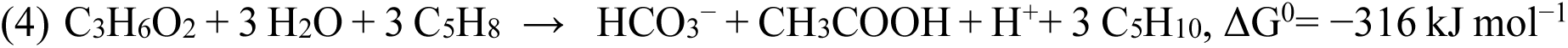

In our study, isoprene was consistently reduced to methyl butenes, and *Pelotomaculum* was the most abundant microorganism in all enrichments supplemented with both isoprene and carbon sources. *Pelotomaculum* was present at a low relative abundance (0.1% of the total microbial community) in cultures without isoprene but became the most dominant species (18% of the total microbial community) in the presence of isoprene (Fig. 2C). In the second-dilution cultures, *Pelotomaculum* accounted for 30-31% of the total microbial community. *Pelotomaculum schinkii* and *Pelotomaculum* sp. FP were the most dominant species enriched throughout the dilution-to-extinction experiment. These findings strongly support our hypothesis that *Pelotomaculum* utilizes isoprene as an alternative electron acceptor, providing a competitive growth advantage.

This hypothesis is further supported by a homology search for *isrA* (a FAD-dependent oxidoreductase) implicated in isoprene reduction in *Acetobacterium wieringae* strain ISORED-2. In *Acetobacterium wieringae* strain ISORED-2, Kronen *et al*. (2023) identified an *isr* operon consisting of 5 genes, including a FAD-dependent oxidoreductase-coding gene (*isrA*), 3 pleiotropic nickel chaperone-coding genes (2 *HypA*, *HypB*), and 4Fe-4S ferredoxin-coding genes. Jin *et al*. (2022) also reported high expression of a FAD-dependent oxidoreductase (LNN31_08025) in isoprene-and HCO ^-^/H_2_-amended cultures of *A. wieringae* strain Y. LNN31_08025 shares 100% nucleotide identity with *isrA*, and the five-gene operon that encodes LNN31_08025 shares 100% nucleotide similarity with the *isr* operon. Kronen *et al*. (2023) found that *isrA* (VUZ27132.1) has the closest amino acid similarity to an FAD-dependent oxidoreductase in *Pelotomaculum* species and reported that *Pelotomaculum schinkii* harbors an FAD-dependent oxidoreductase (WP_190259616.1) with 77.42% amino acid similarity, and *Pelotomaculum* sp.FP harbors an FAD-dependent oxidoreductase (WP_192895435.1) that shares 52% amino acid similarity with protein VUZ27132.1, the putative isoprene reductase in *Acetobacterium wieringae* strain ISORED-2. Other *Acetobacterium* species exhibit significantly lower amino acid similarity (ranging from 47% to 49%) to *isrA*, suggesting that *isrA* is more closely related to *Pelotomaculum* species than to any other *Acetobacterium* species.

*Pelotomaculum schinkii* shows the highest amino acid similarity to *isrA*, yet its *isrA* operon lacks the second *HypA* gene and the ferredoxin gene, rendering the operon incomplete. This raises the question of whether these missing genes are essential for isoprene reduction. Further dilution-to-extinction studies are needed to determine whether isoprene reduction in our enrichment cultures results from the activity of *Pelotomaculum schinkii* alone or from a combined effort among multiple microorganisms.

### Isoprene reduction by *Pelotomaculum* severs the syntrophic association between *Pelotomaculum* and a hydrogenotrophic methanogen

In anaerobic microbial communities, syntrophic interactions between bacteria and methanogens are crucial for methane production. These interactions depend on the transfer of soluble electron carriers, including hydrogen (H₂), formate, and acetate, from bacterial partners to methanogens (Thauer *et al.,* 2008; Yin *et al*., 2020). Because these processes occur near thermodynamic equilibrium, they are highly sensitive to environmental changes (Yin *et al*., 2020).

Species of *Pelotomaculum*, including *P. propionicum* (Imachi *et al*., 2007), *P. schinkii* (de Bok *et al*., 2005), and *P. thermopropionicum* (Imachi, 2002), are known syntrophic anaerobes that oxidize propionic acid, butyric acid, alcohols, phthalate isomers (Kleerebezem *et al*., 1999; Qiu *et al*., 2004, 2006), phenol (Wang *et al*., 2020), and petrochemical waste (D. Wang *et al.,* 2020) to acetate, hydrogen (H₂), and carbon dioxide (CO₂). These end products are then consumed by methanogens to produce methane (Kato *et al*., 2009). Under typical methanogenic conditions, *Pelotomaculum* species generate reducing equivalents, such as NADH, reduced ferredoxin (Fd_red), and menaquinol. To sustain substrate oxidation, these reducing equivalents must be reoxidized.

Typically, H⁺ (E^0^’ = −414 mV) and CO₂ (E^0^’ = −430 mV) serve as electron sinks for *Pelotomaculum*. However, because their reduction potentials are lower than that of NADH (E^0^’ = −320 mV), these reductions are thermodynamically unfavorable (Nobu *et al*., 2015; Hidalgo-Ahumada *et al*., 2018). To overcome this challenge, *Pelotomaculum* uses hydrogenases and formate dehydrogenases to couple the unfavorable oxidation of NADH with the favorable oxidation of Fd_red, thereby driving the reduction of H⁺ and CO₂ to H₂ and formate, respectively (Nobu *et al*., 2015; Hidalgo-Ahumada *et al.,* 2018). To sustain the oxidation of substrates such as propionate, *Pelotomaculum* relies on methanogens to maintain low concentrations of H₂ and formate.

In our microcosm experiments, we observed a decline in the relative abundance of methanogens and methane production over time in all enrichments containing isoprene, with methane production undetectable in dilution cultures. In contrast, isoprene reduction activity remained consistent, and *Pelotomaculum* abundance increased over time across all enrichment and dilution cultures. This suggests that isoprene serves as an alternative electron acceptor for *Pelotomaculum*. Kronen *et al*. (2023) previously suggested that isoprene reduction could be linked to energy conservation in *Pelotomaculum*, though this was not directly measured in our study. The increased relative abundance of *Pelotomaculum* in cultures supplemented with isoprene supports this hypothesis. By acting as an electron sink, isoprene may enable *Pelotomaculum* to oxidize substrates more efficiently, bypassing the need for methanogens to regulate hydrogen and formate concentrations.

Moderate inhibition of methanogenesis by isoprene has been reported previously (Schink, 1985; Kronen *et al*., 2019; Jin *et al*., 2022), yet the precise mechanisms remain unclear. In our study, cultures supplemented with both carbon sources and isoprene showed a significant reduction in methane production compared with those receiving only carbon sources. 16S rRNA analysis revealed distinct microbial dynamics: *Methanosarcina* was the dominant methanogen in the presence of isoprene, whereas *Methanobacterium* predominated in cultures without isoprene. This shift is significant because *Methanobacterium* species rely exclusively on hydrogenotrophic methanogenesis, whereas *Methanosarcina* can utilize hydrogenotrophic, methylotrophic, and acetoclastic methanogenic pathways. This metabolic flexibility likely allows *Methanosarcina* to persist despite reduced hydrogen availability.

The potential ability of *Pelotomaculum* to use isoprene as an electron acceptor likely reduces the hydrogen pool available to methanogens. Although hydrogen-producing bacteria were present in our microcosm, the suppression of *Methanobacterium* in the presence of isoprene suggests competition for hydrogen between methanogens and isoprene reducers. These findings provide novel insights into the microbial interactions governing isoprene and methane metabolism, underscoring the need for further investigation to fully elucidate these dynamics.

## Conclusion

Our study highlights the significant role of *Pelotomaculum* species in isoprene reduction within anaerobic microbial communities. The ability of *Pelotomaculum* to use isoprene as an electron acceptor suggests a novel metabolic pathway that disrupts its syntrophic association with methanogens. Consistent isoprene-reducing activity and increased relative abundance of *Pelotomaculum* in isoprene-amended microcosms expand the known diversity of isoprene-reducing anaerobes beyond *Acetobacterium wieringae*, which was previously the only anaerobic species known to reduce isoprene. The shift in microbial interactions observed in our study, particularly the suppression of hydrogenotrophic methanogens such as *Methanobacterium* and the persistence of *Methanosarcina*, underscores the broader ecological and biogeochemical implications of isoprene reduction. Given these findings, future research should focus on elucidating the specific biochemical mechanisms and genetic pathways involved in isoprene reduction by *Pelotomaculum*. Detailed genomic, proteomic, and enzymatic analyses are essential to identify the key enzymes and regulatory factors driving this process. Additionally, isolating and characterizing *Pelotomaculum* strains with isoprene-reducing capabilities will provide deeper insights into the diversity and evolution of this metabolic trait.

## Supporting information

Figure S4. Scanning electron microscopy (SEM) images showing cellular morphologies and aggregate structures in eucalyptus-derived anaerobic enrichment

Figure S2. Time-course profiles of isoprene, its reduced products; 2-methyl-1-butene and 3-methyl-1-butene (a), and methane concentrations (b) in euca

## Acknowledgements

This work was supported by the College of Arts and Sciences (CAS) at Oklahoma State University (OSU). Support was also provided by the Start-up Grant Package from the Department of Microbiology and Molecular Genetics at Oklahoma State University to SB; the Arts and Sciences Research Award (ARS) to SB; the Roberson Summer Dissertation Fellowship to SG; and the ASM Future Leaders Mentoring Fellowship (ASM_FLMF) to SG. Sequence analyses were carried out at the Microbial Analysis, Resources, and Services Center for Open Research Resources and Equipment at the University of Connecticut and at BMKGENE. SEM microscopy was performed at the Oklahoma State University Microscopy Facility with the help of Lisa Withworth. Infrastructural support was provided by Dr. Roger Mailler in the Department of Computer Sciences.

## Author Contributions

Samikshya Giri and Dr. Sabrina Beckmann contributed to the design and implementation of the research, analysis of the results, and manuscript writing. Dr. Sabrina Beckmann conceived the original idea and supervised the project. Anna E. Rockwood and Zoey Ilene Kelley assisted with experimental work and data collection.

## Supplementary file

**Figure S1.**
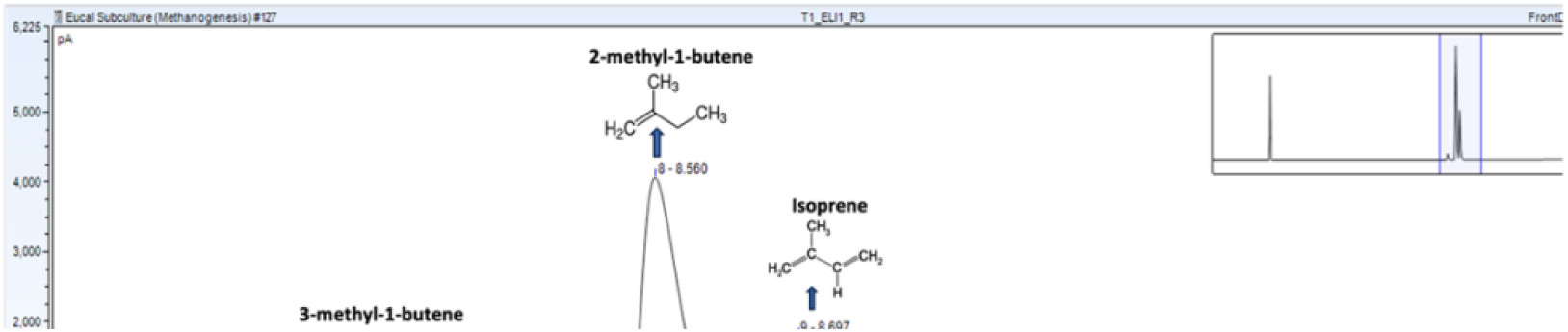
Representative gas chromatography–flame ionization detector (GC-FID) chromatogram from an active enrichment culture showing isoprene and its reduced products. Peaks corresponding to isoprene, 2-methyl-1-butene, 3-methyl-1-butene, and 2-methyl-2-butene are indicated based on retention time and comparison with authentic standards.

**Figure S2.**
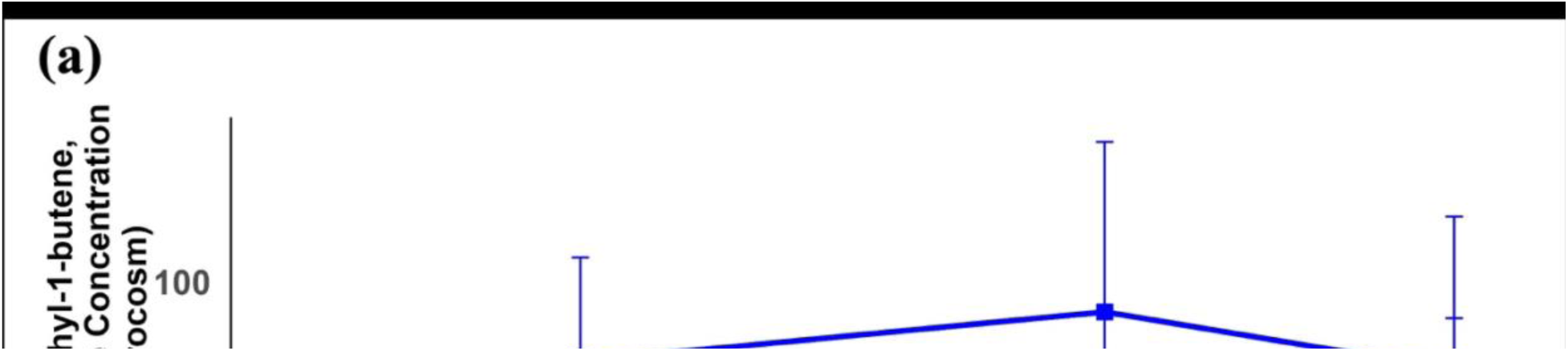
Time-course profiles of isoprene, its reduced products; 2-methyl-1-butene and 3-methyl-1-butene (a), and methane concentrations (b) in eucalyptus-derived sediment enrichment cultures incubated with or without isoprene amendment. Measurements were collected from week 1 following re-addition of fresh media and isoprene through 8 weeks of incubation.

**Figure S3.**
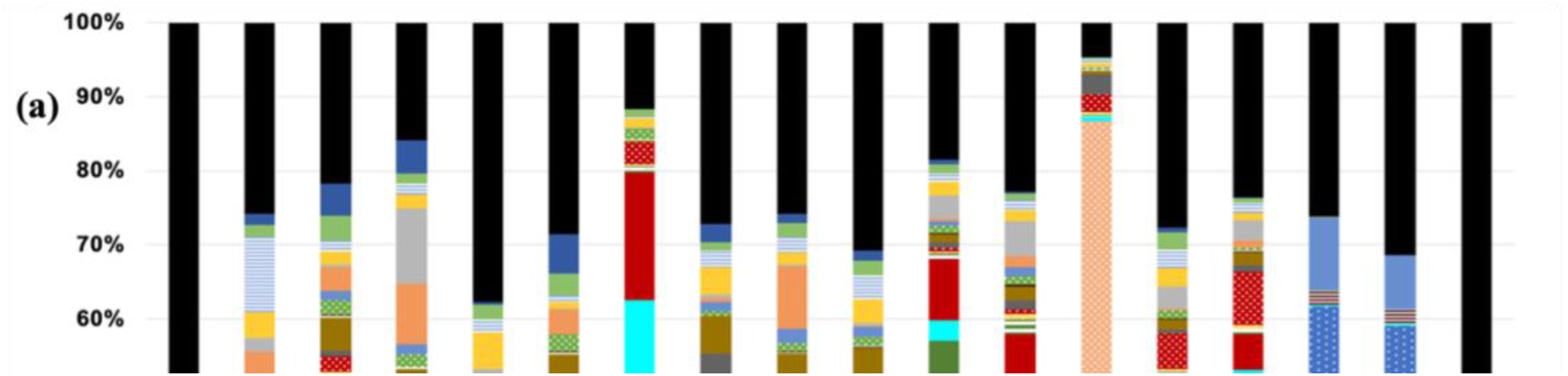
(a) Total Relative abundance of microbial taxa based on 16S rRNA gene sequencing in eucalyptus-leaf sediment microcosms amended with volatile fatty acids (VFAs) and lactate, with and without isoprene, and the dead control at 3 different time points (T1: 2 months incubation, T2: 12 months incubation, Re_T3: 1 week after the re-addition of media, substrate, and isoprene). The original soil inoculum at Time 0 is labeled as (ORI_T0). Stacked bar plots show the proportional abundance of dominant taxa across individual microcosms. Taxa below the defined relative (<1%) abundance threshold are grouped as “Others.” **(b)** Total relative abundance of bacterial taxa inferred from 16S rRNA gene sequencing in eucalyptus-leaf sediment microcosms incubated with and without isoprene and the dead control at 3 different time points (T1: 2 months incubation, T2: 12 months incubation, Re_T3: 1 week after the re-addition of media, substrate, and isoprene). Original soil inoculum at Time 0 is labelled as ORI_T0. Community composition is shown for individual replicates and enrichment transfers across three time points (T1: 4 months, T2: 12 months, Re_T3: 1 weeks after the replenishment of media and isoprene). Only taxa exceeding a defined relative abundance threshold of 0.1% are displayed individually; remaining taxa are grouped as “Others.”

**Figure S4.**
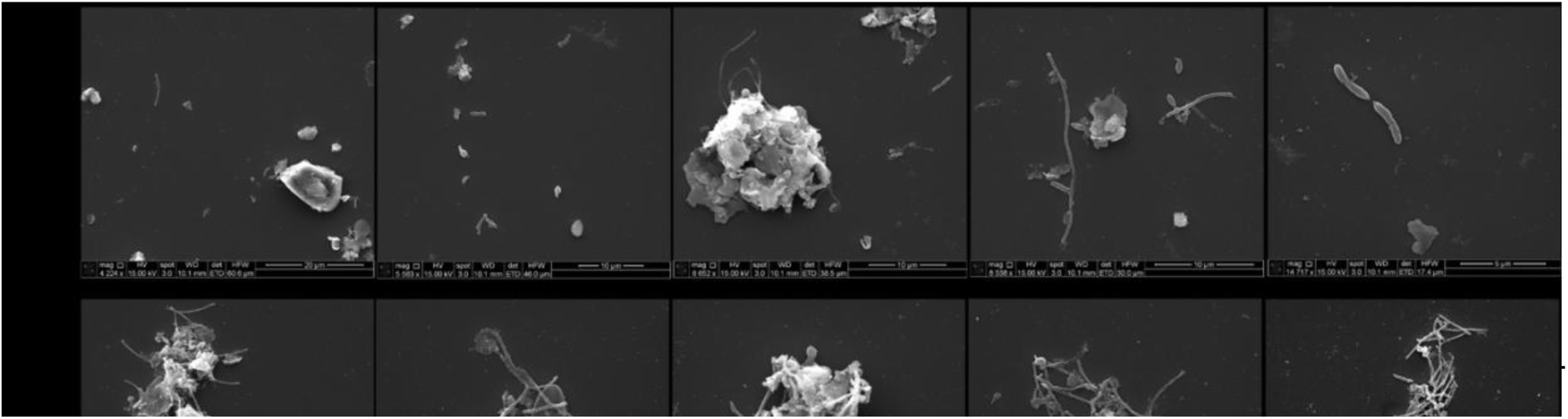
Scanning electron microscopy (SEM) images showing cellular morphologies and aggregate structures in eucalyptus-derived anaerobic enrichment cultures amended with isoprene, lactate, and volatile fatty acids (VFAs). **(a)** Original enrichment culture (Eucal + Isoprene + Lactate + VFAs_1). **(b)** First dilution culture derived from the original enrichment. **(c)** Subsequent dilution culture derived from the same enrichment. Images illustrate the diversity of cell morphologies and extracellular structures observed across enrichment and dilution steps.

**Figure S5.**
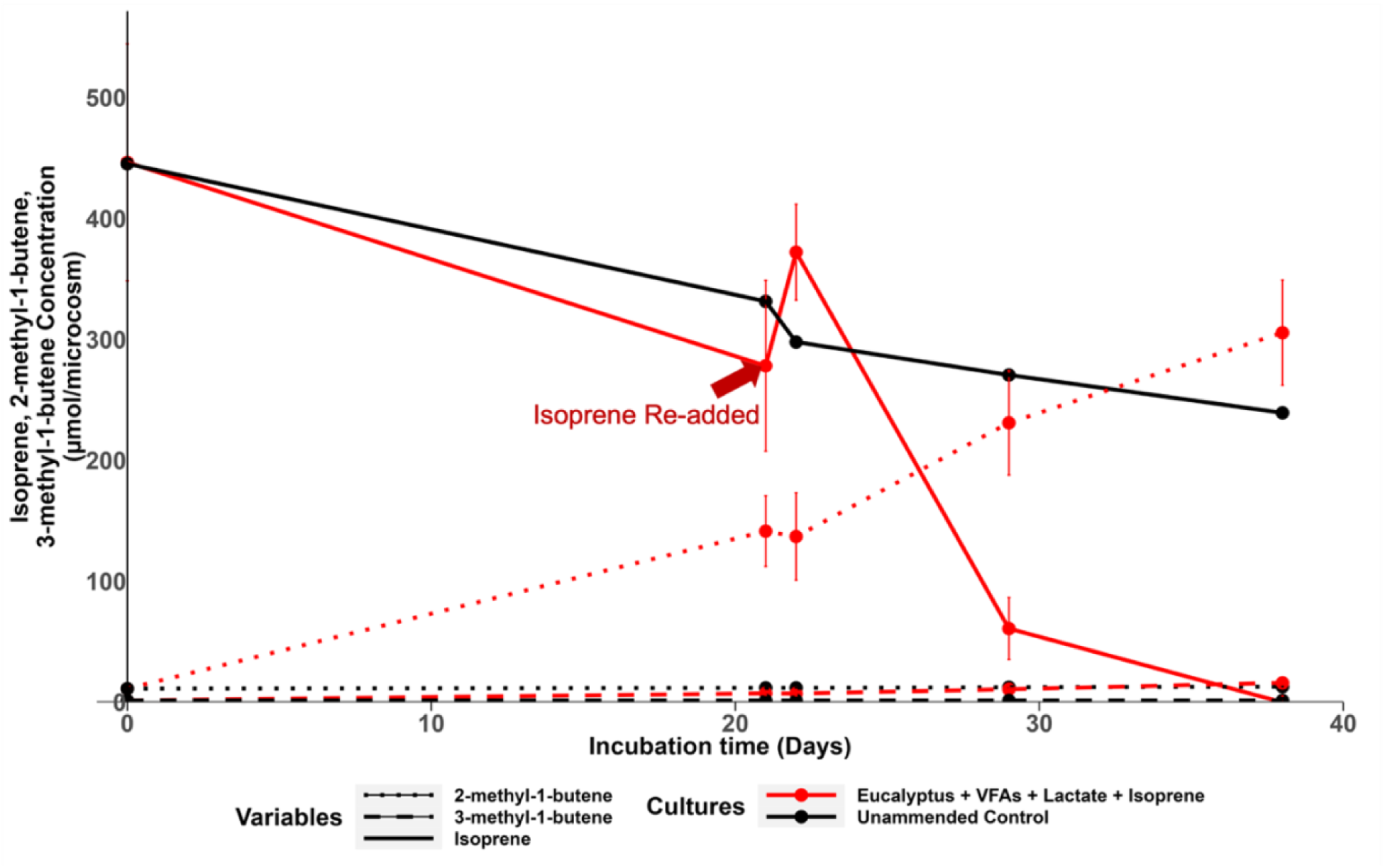
Time-course of isoprene and reduced isoprene products in the sixth-transfer enrichment culture ICE1. Concentrations of isoprene, 2-methyl-1-butene, and 3-methyl-1-butene were monitored in cultures amended with inoculum volatile fatty acids, lactate, and isoprene (red) and in unamended controls (black, no inoculum). Isoprene re-addition during incubation is indicated by the arrow. Data points represent measured concentrations, and error bars indicate variability among biological replicates (n = 4), where applicable.

## References

Ashworth K, Wild O, & Hewitt CN (2013) Impacts of biofuel cultivation on mortality and crop yields. Nature Climate Change, 3(5), 492–496. 10.1038/nclimate1788

Beckmann S & Manefield M (2014) Acetoclastic methane formation from eucalyptus detritus in pristine hydrocarbon-rich river sediments by Methanosarcinales. FEMS Microbiology Ecology, 90(3), 587–598. 10.1111/1574-6941.12418

Caporaso JG, Lauber CL, Walters WA, Berg-Lyons D, Lozupone CA, Turnbaugh PJ et al. (2011) Global patterns of 16S rRNA diversity at a depth of millions of sequences per sample. PNAS, 108(Suppl 1), 4516–4522. 10.1073/pnas.1000080107

Carrión O, McGenity TJ, & Murrell JC (2020) Molecular ecology of isoprene-degrading bacteria. Microorganisms, 8(7), 967. 10.3390/microorganisms8070967

Charlson RJ, Schwartz SE, Hales J, Cess RD, Coakley JA, Hansen JE et al. (1992) Climate forcing by anthropogenic aerosols. Science, 255(5043), 423–430. 10.1126/science.255.5043.423

Cleveland CC & Yavitt JB (1997) Consumption of atmospheric isoprene in soil. Geophysical Research Letters, 24(19), 2379–2382. 10.1029/97gl02451

Collins WJ, Derwent RG, Johnson CE & Stevenson DS (2002) Climatic Change, 52(4), 453–479. 10.1023/a:1014221225434

Crombie AT, Khawand ME, Rhodius VA, Fengler KA, Miller MC, Whited GM et al. (2015) Regulation of plasmid-encoded isoprene metabolism in *Rhodococcus*. Environmental Microbiology, 17(9), 3314–3329. 10.1111/1462-2920.12793

Dalton H & Stirling DI (1982) Co-metabolism. Philosophical Transactions of the Royal Society B, 297(1088), 481–496. 10.1098/rstb.1982.0056

Dawson RA, Crombie AT, Jansen RS, Smith TJ, Nichol T & Murrell C (2023) Peering down the sink: A review of isoprene metabolism by bacteria. Environmental Microbiology, 25(4), 786–799. 10.1111/1462-2920.16325

De Bok FAM, Harmsen HJM, Plugge CM, De Vries MC, Akkermans ADL, De Vos WM et al. (2005) *Pelotomaculum schinkii* sp. nov. IJSEM, 55(4), 1697–1703. 10.1099/ijs.0.02880-0

Diez-Gonzalez F, Russell JB & Hunter JB (1995) NAD-independent lactate dehydrogenase in *Clostridium acetobutylicum*. Archives of Microbiology, 164(1), 36–42. 10.1007/s002030050233

Du Y, Zou W, Zhang K, Ye G & Yang J et al. (2020) Advances and applications of *Clostridium* co-culture systems. Frontiers in Microbiology, 11. 10.3389/fmicb.2020.560223

Ervens B, Turpin BJ & Weber RJ (2011) Secondary organic aerosol formation in cloud droplets. Atmospheric Chemistry and Physics, 11(21), 11069–11102. 10.5194/acp-11-11069-2011

Fall R & Copley SD (2000) Bacterial sources and sinks of isoprene. Environmental Microbiology, 2(2), 123–130. 10.1046/j.1462-2920.2000.00095.x

García-Depraect O, Castro-Muñoz R, Muñoz R, Rene ER, León-Becerril E, Valdez-Vazquez I et al. (2021) Factors influencing biohydrogen production from lactate. Bioresource Technology, 324, 124595. 10.1016/j.biortech.2020.124595

Gelmont D, Stein RA & Mead JF (1981) Isoprene in human breath. Biochemical and Biophysical Research Communications, 99(4), 1456–1460. 10.1016/0006-291x(81)90782-8

Gray CM, Helmig D & Fierer N (2015) Isoprene consumption in soil. Elementa, 3. 10.12952/journal.elementa.000053

Hidalgo-Ahumada CP, Nobu MK, Narihiro T, Tamaki H, Liu W, Kamagata Y et al. (2018) Energy conservation strategies in *Pelotomaculum schinkii*. Environmental Microbiology, 20(12), 4503–4511. 10.1111/1462-2920.14388

Hoang VT, Hoang DH, Pham ND, Tran HM, Bui HTV & Ngo TA (2018) Hydrogen production by newly isolated Clostridium species from cow rumen in pure and co-cultures on a broad range of carbon sources. AIMS Energy, 6(5), 846–865. 10.3934/energy.2018.5.846

Imachi H, Sekiguchi Y, Kamagata Y, Hanada S, Ohashi A & Harada H (2002) *Pelotomaculum thermopropionicum* gen. nov., sp. nov. IJSEM, 52(5), 1729–1735. 10.1099/00207713-52-5-1729

Imachi H, Sakai S, Ohashi A, Harada H, Hanada S, Kamagata Y et al. (2007) *Pelotomaculum propionicicum* sp. nov. IJSEM, 57(7), 1487–1492. 10.1099/ijs.0.64925-0

Jin H, Li X, Wang H, Cápiro NL, Li X, Löffler FE et al. (2022) Anaerobic biohydrogenation of isoprene by *Acetobacterium wieringae*. mBio, 13(6). 10.1128/mbio.02086-22

Kato S, Kosaka T & Watanabe K (2009) Substrate-dependent transcriptomic shifts in *Pelotomaculum thermopropionicum grown in syntrophic co-culture with Methanothermobacter thermautotrophicus*. Microbial Biotechnology, 2(5), 575–584. 10.1111/j.1751-7915.2009.00102.x

Khawand ME, Crombie AT, Johnston A, Vavlline DV, McAuliffe JC et al. (2016) Isolation of isoprene-degrading bacteria and isoA probes. Environmental Microbiology, 18(8), 2743–2753. 10.1111/1462-2920.13345

Kleerebezem R, Pol LWH & Lettinga G (1999) Anaerobic degradation of phthalate isomers by Methanogenic Consortia. Applied and Environmental Microbiology, 65(3), 1152–1160. 10.1128/AEM.65.3.1152-1160.1999

Kronen M, Lee M, Jones ZL & Manefield MJ (2019) Reductive metabolism of the important atmospheric gas isoprene by homoacetogens. ISME Journal, 13(5), 1168–1182. 10.1038/s41396-018-0338-z

Kronen M, Vázquez-Campos X, Wilkins MR, Lee M & Manefield MJ (2023) Evidence for a Putative Isoprene Reductase in *Acetobacterium wieringae*. mSystems, 8(2). 10.1128/msystems.00119-23

Lawson PA, Wawrik B, Allen TD, Johnson CN, Marks CR, Tanner RS et al. (2014) Youngiibacter fragilis gen. nov., sp. nov., isolated from natural gas production-water and reclassification of Acetivibrio multivorans as Youngiibacter multivorans comb. nov. IJSEM, 64(Pt 1), 198–205. 10.1099/ijs.0.053728-0

Lunau M, Lemke A, Walther K, Martens-Habbena W & Simon M (2005) An improved method for counting bacteria from sediments and turbid environments by epifluorescence microscopy. Environmental Microbiology, 7(7), 961–968. 10.1111/j.1462-2920.2005.00767.x

McGenity TJ, Crombie AT & Murrell JC (2018) Microbial cycling of isoprene, the most abundantly produced biological volatile organic compound on Earth. ISME Journal, 12(4), 931–941. 10.1038/s41396-018-0072-6

Murrell JC, McGenity TJ, & Crombie AT (2020) Microbial metabolism of isoprene: a much-neglected climate-active gas. Microbiology Society, 166(7), 600–613. 10.1099/mic.0.000931

Nobu MK, Narihiro T, Rinke C, Kamagata Y, Tringe SG, Woyke T et al. (2015) Microbial dark matter ecogenomics reveals complex synergistic networks in a methanogenic bioreactor. ISME Journal, 9(8), 1710–1722. 10.1038/ismej.2014.256

Pacifico F, Harrison S, Jones C & Sitch S (2009) Isoprene emissions and climate. Atmospheric Environment, 43(39), 6121–6135. 10.1016/j.atmosenv.2009.09.002

Parada AE, Needham DM & Fuhrman, JA (2016) Every base matters: assessing small subunit rRNA primers for marine microbiomes with mock communities, time series and global field samples. Environmental Microbiology, 18(5), 1403–1414. 10.1111/1462-2920.13023

Qiu Y, Sekiguchi Y, Hanada S, Imachi H, Tseng I, Cheng S et al. (2006) *Pelotomaculum terephthalicum* sp. nov. *and Pelotomaculum isophthalicum* sp. nov.: two anaerobic bacteria that degrade phthalate isomers in syntrophic association with hydrogenotrophic methanogens. Arch Microbiol 185, 172–182 (2006). 10.1007/s00203-005-0081-5

Schink B (1985) Inhibition of methanogenesis by ethylene and other unsaturated hydrocarbons, FEMS Microbiology Ecology, Volume 1, Issue 2, April 1985, Pages 63–68. 10.1111/j.1574-6968.1985.tb01132.x

Schloss PD, Westcott SL, Ryabin T, Hall JR, Hartmann M, Hollister EB et al. (2009) Introducing mothur: Open-Source, Platform-Independent, Community-Supported Software for Describing and Comparing Microbial Communities. Applied and Environmental Microbiology, 75, 7537–7541. 10.1128/AEM.01541-09

Sharkey TD, Wiberley AE & Donohue AR (2008) Isoprene Emission from Plants: Why and How, Annals of Botany, Volume 101, Issue 1, January 2008, Pages 5–18. 10.1093/aob/mcm240

Singh A, Schnürer A, Dolfing J & Westerholm M (2023) Syntrophic entanglements for propionate and acetate oxidation under thermophilic and high-ammonia conditions. The ISME Journal, Volume 17, Issue 11, November 2023, Pages 1966–1978. 10.1038/s41396-023-01504-y

Sims LP, Lockwood CWJ, Crombie AT, Bradley JM, Le Brun NE, Murrell JC et al. (2022) Purification and Characterization of the Isoprene Monooxygenase from *Rhodococcus* sp. Strain AD45. Appl Environ Microbiol 88:e00029-22. 10.1128/aem.00029-22

Thauer RK, Kaster AK, Seedorf H, Buckel W, & Hedderich R (2008) Methanogenic archaea: ecologically relevant differences in energy conservation. Nat Rev Microbiol 6, 579–591 (2008). 10.1038/nrmicro1931

Wang J, Wu B, Sierra JM, He C, Hu Z, & Wang W (2020) Influence of particle size distribution on anaerobic degradation of phenol and analysis of methanogenic microbial community. Environ Sci Pollut Res 27, 10391–10403 (2020). 10.1007/s11356-020-07665-z

Wang J & Yin Y (2021) *Clostridium* species for fermentative hydrogen production: An overview. International Journal of Hydrogen Energy, 46(70), 34599–34625. 10.1016/j.ijhydene.2021.08.052

Wang D, Gao C, Wang C, Liu N, Qiu C, Yu J, & Wang S (2021) Effect of mixed petrochemical wastewater with different effluent sources on anaerobic treatment: organic removal behaviors and microbial community. Environ Sci Pollut Res 28, 5880–5891 (2021). 10.1007/s11356-020-10951-5

Westerholm M, Calusinska M & Dolfing J (2022) Syntrophic propionate-oxidizing bacteria in methanogenic systems, *FEMS Microbiology Reviews*, Volume 46, Issue 2, March 2022, fuab057, 10.1093/femsre/fuab057

Widdel F & Bak F (1992) Gram-Negative Mesophilic Sulfate-Reducing Bacteria. In: Balows, A., Trüper, H.G., Dworkin, M., Harder, W., Schleifer, KH. (eds) The Prokaryotes. Springer, New York, NY. 10.1007/978-1-4757-2191-1_21.

Xue J & Ahring BK (2011) Enhancing Isoprene Production by Genetic Modification of the 1-Deoxy-d-Xylulose-5-Phosphate Pathway in Bacillus subtilis. Appl Environ Microbiol 77: 10.1128/AEM.02341-10

Yin Q, Gu M & Wu G (2020) Inhibition mitigation of methanogenesis processes by conductive materials: A critical review. Bioresource Technology, 317. 10.1016/j.biortech.2020.123977

