## Supplementary figures and images for "Anaerobic isoprene reduction by *Pelotomaculum* sp. From *Eucalyptus*-leaf sediments and its impacts on methanogenesis"

### Figure S2. Time-course profiles of isoprene, its reduced products; 2-methyl-1-butene and 3-methyl-1-butene (a), and methane concentrations (b) in euca

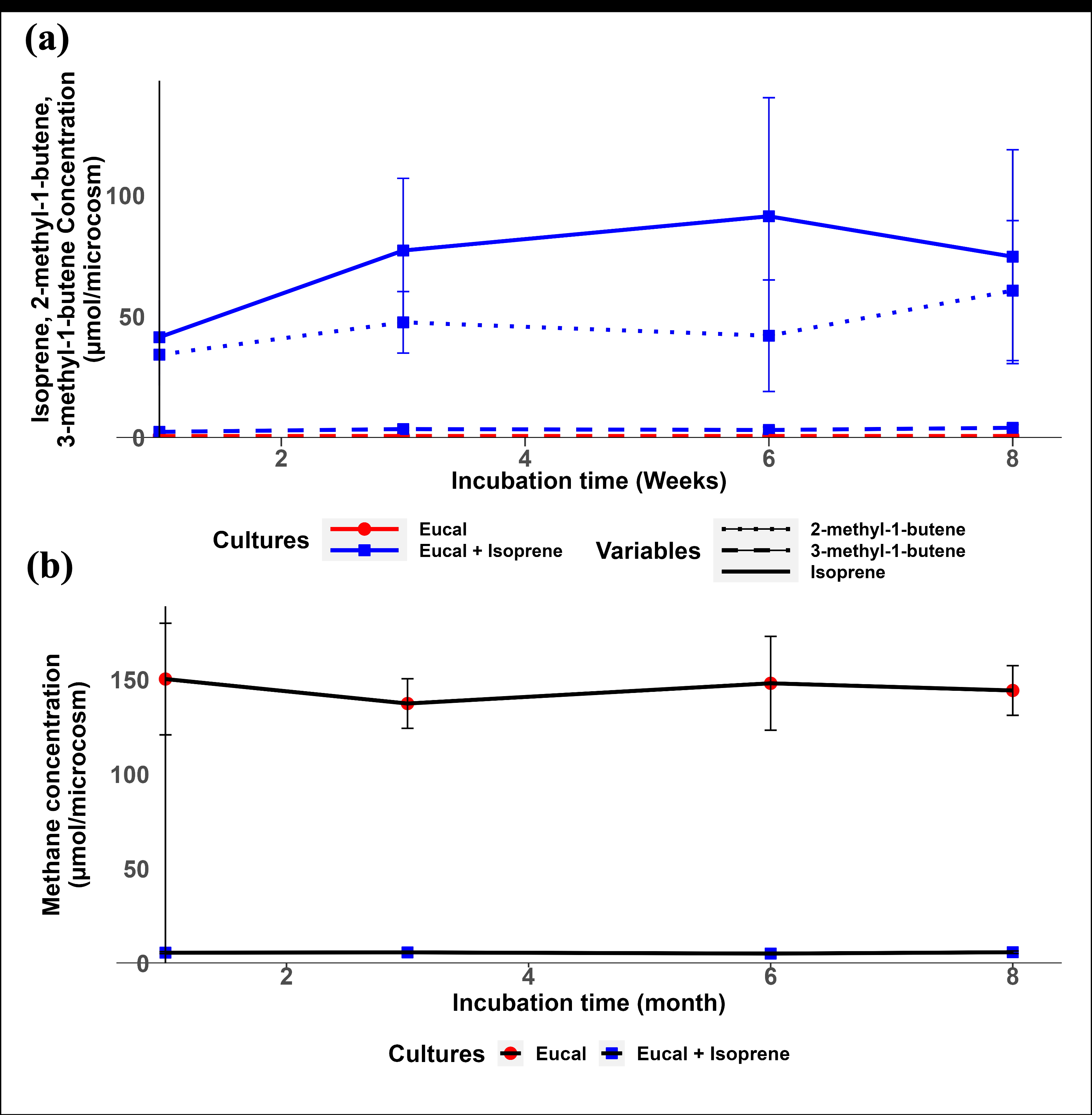

### Figure S4. Scanning electron microscopy (SEM) images showing cellular morphologies and aggregate structures in eucalyptus-derived anaerobic enrichment

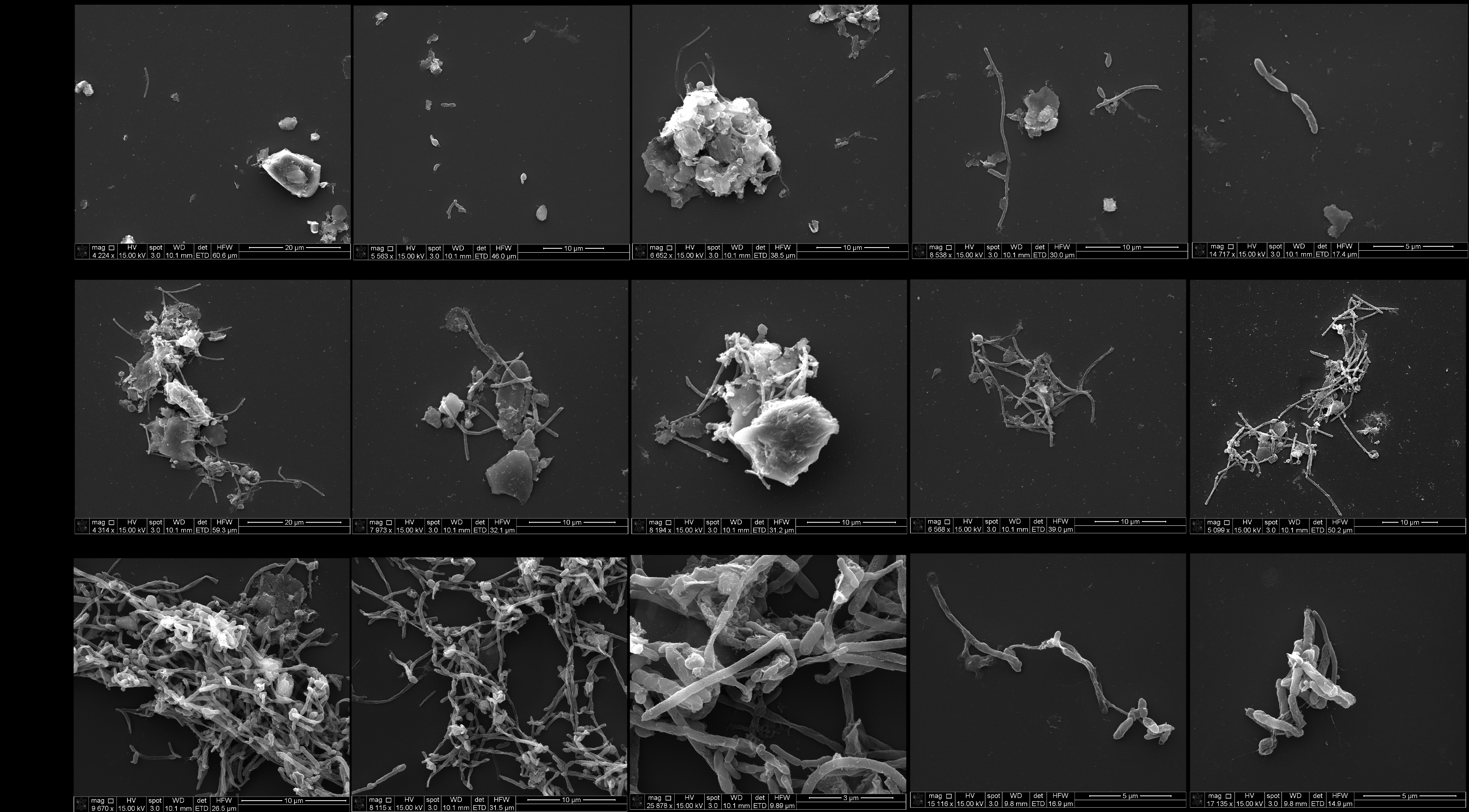
